# Red and blue light cues drive contrasting remodeling of lipophilic metabolites and photophysiology in natural benthic diatom biofilms

**DOI:** 10.64898/2026.08.30.748109

**Authors:** Alexandre Desparmet, Johann Lavaud, Bruno Jesus, Antoine Medico, Cédric Hubas

## Abstract

Intertidal mudflats are low hydrodynamic energy environments hosting microphytobenthic communities that experience strong spatiotemporal variability in light regimes, including changes in spectral quality and light intensity that can lead to cellular photooxidative stress. To cope with these fluctuations, autotrophs exhibit diverse and highly plastic adaptations that are often species-dependent and shaped by their ecological niches. This study investigates photophysiological responses and metabolic remodeling in a diatom assemblage originating from a natural winter microphytobenthic biofilm under contrasting red and blue light intensities. To this end, photosynthetic parameters were monitored alongside changes in lipophilic metabolites, including untargeted lipids and lipophilic pigments. While few metabolites showed temporal remodeling, rapid and contrasting changes were observed within 30 minutes in response to both spectral quality and light intensity. Red light treatments induced broader remodeling of lipophilic metabolites than blue light, whereas blue light appeared to have a greater impact on photosynthetic parameters. Moreover, red light induced xanthophyll-cycle responses comparable to those observed under blue light at equivalent incident intensity. We discuss these metabolic responses in relation to diatom photoadaptive strategies, placing these findings within the intertidal environmental framework. This work further underlines the importance of understanding rapid metabolic plasticity in coping with light fluctuations, providing new insights into the photoregulatory strategies of natural microphytobenthic communities.

## Introduction

Microphytobenthic biofilms inhabit cohesive intertidal mudflat sediments and form three-dimensional, mucilaginous, surface-attached communities characterized by high microalgal biomass, with epipelic diatoms frequently being the dominant group (Underwood & Barnett, 2006; Hubas et al., 2018; Desparmet et al., 2026). Tidal environments undergo strong spatial and temporal fluctuations of numerous physical and chemical environmental factors. Among these, light intensity and spectral composition can vary depending on diurnal, tidal, and seasonal cycles (Perkins et al., 2001; Passarelli et al., 2015; Underwood et al., 2022), as well as on interactions between light and the water column (e.g., turbidity, depth) (Depauw et al., 2012; Brunet et al., 2014) and the sediment (e.g., granulometry, organic matter, pigment community composition) (Kühl and Jørgensen, 1994; Consalvey et al., 2004; Jesus et al. 2009; Cartaxana et al., 2011). Within a few minutes, light conditions experienced by microphytobenthos can shift rapidly to over 2000 µmol photons m^-2^ s^-1^ (Waring et al., 2010), disrupting linear electron flow and efficient energy transfer from photosystem II (PSII) to photosystem I (PSI) (Shimakawa and Miyake, 2018; Xu et al., 2020; Lepetit et al., 2022). These conditions lead to an over-stimulation of the photosynthetic electron transport chain, whereby the capacity to dissipate excess absorbed light energy becomes saturated, while continued light absorption promotes photooxidative damage and, when cellular antioxidant capacity is exceeded, photoinhibition (Serôdio et al., 2012; Foyer et al., 2022). To maintain photosynthetic efficiency throughout the day, diatoms use several physiological responses enabling adaptation to rapid light changes.

One of the major mechanisms is the use of non-photochemical quenching (NPQ) mediated by the xanthophyll-cycle, enzymatically regulated via lumen acidification induced by the trans-thylakoid proton gradient under high light (Blommaert, 2017; Goss & Latowski, 2020; Lepetit et al., 2022). The conversion of diadinoxanthin to diatoxanthin allows excess energy to be dissipated as heat, thereby limiting the increase in oxidative stress and protecting photosystems integrity (Foyer et al., 2022; Lepetit et al., 2022). This rapid photoregulatory process (Blommaert et al., 2017; Laviale et al., 2016) acts synergistically with vertical migration within sediments, allowing benthic diatoms to quickly position themselves along light gradients (Cartaxana et al., 2011; Jesus et al., 2023; Morelle et al., 2024; Desparmet et al., 2026).

Moreover, additional processes occur in response to spectral and intensity changes, including the reorganization of the partitioning of photosynthetically assimilated carbon, particularly within the lipid macromolecular pool, which represents approximately 20% of diatom dry weight (Finkel et al., 2016; Wagner et al., 2017). Rapid cellular adjustments are regulated by both redox homeostasis, notably shaped by light-dependant reactive oxygen species production (Waring et al., 2010; Foyer et al., 2022; Doose & Hubas, 2024), and by direct photoreceptor-mediated sensing, including aureochromes, cryptochromes, and rhodopsins sensitive to blue and/or green wavelengths, whereas phytochromes integrate light signals across the visible spectrum, with particularly high sensitivity to red wavelengths (Depauw et al., 2012; Lepetit and Dietzel, 2015; Poliner et al., 2022; Duchêne et al., 2026; Font-Muñoz et al., 2026). Rapid shifts between red and blue light are known to trigger major changes in metabolome and macromolecular composition (Jungandreas et al., 2014), with an interplay between spectral composition and intensity required to drive effective photoacclimation responses (Schellenberger Costa et al., 2013; Brunet et al., 2014; Svenning et al., 2024).

Constantly challenged by their natural environment, the adjustments observed in diatoms are strongly species-specific and partly shaped by previous spectral and intensity light history, aiming to optimize growth and photosynthetic fitness within a specific ecological niche (Van Leeuwe et al., 2008; Croteau et al., 2021; Duarte et al., 2021; Croteau et al., 2022). High metabolic plasticity and sophisticated cellular sensing systems are considered key drivers of the ecological success of diatoms and their dominance in marine autotrophic communities, enabling cells to detect fluctuating conditions and respond adequately (Falciatore et al., 2000; Falciatore et al., 2020; Pierella Karlusich et al., 2024).

Nevertheless, studies investigating metabolic responses at the scale of natural communities remain limited. This study aimed to test the response of a diatom assemblage originating from a natural winter microphytobenthic community to contrasting light intensity regimes under blue or red monochromatic wavelengths. The working hypothesis of distinct photoregulatory responses under these treatments was assessed through the analysis of lipophilic metabolite remodeling, including untargeted lipids and lipophilic pigment, alongside photosynthetic parameters.

## Materials and methods

### Biofilm sampling and diatom suspension

The upper centimeter of sediment containing a dense natural winter microphytobenthic biofilm was collected using a shovel from an outdoor pond. This pond is subject to a semi-diurnal tidal cycle and retains at least 5 cm of clear water above the muddy surface at low tide, ensuring continuous immersion of the community (Concarneau Marine Station, France; 47°52ʹ05.85ʺN, 3°55ʹ00.51ʺW; 13 January 2025, 10:45). Following transport to the laboratory and 24 hours settling to allow biofilm reconstitution, a diatom suspension was prepared the day prior to the experiment using the lens tissue method (Eaton and Moss, 1966). Cells were recovered by gently shaking the tissues into filtered seawater from the sampling site. This protocol selectively collects motile diatoms while minimizing the inclusion of other microorganisms and removing the sediment matrix to prevent vertical migration, thereby ensuring controlled exposure to light treatments.

For each light treatment, five replicates of 5.5-cm-diameter Petri dishes (*n* = 5) were used, each containing 8 mL of diatom suspension for metabolic analyses; three additional replicates (*n* = 3) were used for photophysiological measurements (Table 1). Cell concentration was determined by triplicate counts of a 1:10 diluted preparation using a 1 mL Sedgewick-Rafter chamber, and taxonomic identification was based on previous metabarcoding characterization (Desparmet et al., 2026).

**Table 1.** Experimental setup. Summary of the materials and treatments tested in this study. Raw incident photosynthetic photon flux density (PPFD, in µmol photons m^-2^ s^-1^).

| Treatments | Dark-adapted control | Blue light |  | Red light |  |
| --- | --- | --- | --- | --- | --- |
| Description | Negative control<br>($< 1$ PPFD) | Low light<br>(200 PPFD) | High light<br>(800 PPFD) | Low light<br>(200 PPFD) | High light<br>(800 PPFD) |
| Time exposure | - | 30 min |  |  |  |
| Experimental materials | Leave in darkness | LED panels (SN-SL3500-342, Photon Systems Instruments) |  |  |  |
| Molecular analyses | n = 5 |  |  |  |  |
| Photophysiological measurements | n = 3 |  |  |  |  |

### Light treatments

Two monochromatic light spectrums were tested (Supplementary Fig. 1A). Red light (570– 680nm, peak at 640 nm) was applied in the morning (15 January 2025), while blue light (410– 500 nm, peak at 450 nm) was applied in the afternoon of the same day. Both light spectra were tested at two intensities: low light (200 µmol photons m^-2^ s^-1^) and high light (800 µmol photons m^-2^ s^-1^). These intensities were selected based on published values (Prins et al., 2020; Goessling et al., 2016; Smerilli et al., 2017), were consistent with preliminary measurements conducted on this natural winter microphytobenthic biofilm prior to the experiment (Supplementary Fig. 1B) and remained below light levels that diatoms may encounter and tolerate under natural conditions (Waring et al., 2010; Desparmet et al., 2026). The intensities indicated above correspond to incident light; additional calculations for estimating package-corrected photosynthetically usable incident light (Qphar_package-corrected_) and the resulting absorbed photon flux and absorbed energy are provided in the Supplementary Materials and Methods (Supplementary Fig. 8).

Samples were exposed to one of the four distinct light treatments (30 minutes each) and compared to dark-adapted controls maintained in room darkness (<1 µmol photons m^-2^ s^-1^, which is not complete darkness, but remains below the light compensation point for microphytobenthos; Walpersdorf et al., 2017). Illumination was provided by LED panels (SN-SL3500-342, Photon Systems Instruments). Light intensity and spectrum were monitored using a MSC15 radiometer (Device-SN 43919, Firmware v1.40, Gigahertz-Optik), coupled with S-MSC15 software v2018.2.1 (DLL v2018.3). Treatments were applied on the same day between 8:30 AM and 12:30 PM for red light, and between 12:00 PM and 4:00 PM for blue light. After the light treatments, the Petri dishes intended for metabolic analyses were immediately frozen in liquid nitrogen, lyophilized, and the resulting dry organic matter was retrieved by gently scraping the dishes before storage at −80 °C.

### Chlorophyll fluorescence: Pulse amplitude modulated fluorometry

Photosynthetic parameters derived from chlorophyll *a* fluorescence were measured using a MonitoringPen MP 100-E fluorometer (Photon System Instruments) to assess photosynthetic performance and interpret subsequent photoprotective responses. Measurements were conducted following a rapid light curve protocol consisting of seven 1-minute intensity steps (λ _LED excitation_ = 470 nm), increasing from 10 to 1000 µmol photons m^-2^ s^-1^ (PPFD), with gain, superpulse, and flashpulse settings maintained consistently throughout all measurements. Relative electron transport rates (rETR) of PSII, reflecting photosynthetic productivity (Perkins et al., 2010), correspond to the values extracted directly from light curve points. The initial linear α-slope, representing the maximum light-use efficiency of photosystem II, was derived from rETR-irradiance (rETR-E) curves using a previously described and modified model (Eilers and Peeters, 1988; Silsbe and Kromkamp 2012). Maximum PSII quantum efficiency of dark-adapted cells (Fv/Fm), together with NPQ and its regulated (Y(NPQ)) and non-regulated (Y(NO)) components, were calculated using established equations (Consalvey et al., 2005; Christof Klughammer & Ulrich Schreiber, 2008) (Supplementary Table 1). Photosynthetic parameters were measured twice for each sample: once before light treatment and once after, following a 10-minute dark recovery period. Dark-adapted control samples were measured alongside experimental samples to ensure identical timing conditions.

### Pigment and lipid analysis

#### Lipophilic pigments processed by HPLC

Pigments were analyzed following the lipophilic pigment extraction protocol from Brotas and Plante-Cuny (2003). Briefly, 50 mg of freeze-dried samples were sonicated for 30 s in 2 mL of 95% methanol buffered with 2% ammonium acetate, extracts were then incubated for 15 min in the dark at −21 °C and filtered through a 0.2 µm PTFE membrane filter. Volumes of 100 µL were injected into an Agilent 1260 Infinity HPLC system equipped with a Supelcosil C18 reverse-phase column. Pigments were separated at a flow rate of 0.6 mL min⁻¹ using solvent A (0.5 M ammonium acetate in 85:15 methanol:water), solvent B (90:10 acetonitrile:water), and solvent C (100% ethyl acetate). Eluted metabolites were identified based on their retention times and absorption spectra recorded with a UV–VIS photodiode array detector (DAD 1260 VL, 250-900 nm). Pigment concentrations (µg of pigment g^-1^ dry weight) were quantified using calibration curves based on phytoplankton pigment standards from Desparmet et al. 2026. For each identified pigment, its relative proportion (as a percentage of the total pigment pool) and its normalized concentration relative to the Σ _[chlorophyll *a*]_ pool (Σ _[chlorophyll *a*]_ = [chlorophyll *a*] + [chlorophyll *a*-derivatives] + [pheopigments]), used as a proxy for biomass, were calculated. In addition, the diadinoxanthin de-epoxidation state (%) was calculated for each light treatment (*n* = 5), as the ratio of diatoxanthin to the total diadinoxanthin+diatoxanthin pool, according to the formula: [diatoxanthin]/([diadinoxanthin] + [diatoxanthin]) x100.

#### Untargeted chloroform-extracted lipids by GC-MS

Lipid profiles from the chloroform phase were analyzed through a simplified version of the untargeted metabolite extraction method used for microphytobenthic samples by Gaubert-Boussarie et al. (2020), and adapted with the Bligh and Dyer (1959) protocol. A 200 mg of freeze-dried mass was extracted three times with 3 mL of a Methanol/Chloroform (MeOH/CHCl_3_, 1:1) mixture in an ultrasonic bath for 30 min with ice packs to promote cell lysis and metabolite desorption. After each extraction step, the mixture was centrifuged for 10 min at 1800g to pellet the dry matter, and the MeOH/CHCl_3_ supernatant was collected. The three extracts were pooled and evaporated under a N_2_ stream. The crude extracts were then resuspended in 4 mL of MeOH/CHCl_3_/H_2_O_Milli-Q_ (2:1:1), vortexed for 20 s, and sonicated for 10 min with ice packs. To complete the biphasic system formed, 2 mL of CHCl_3_/H_2_O_Milli-Q_ (1:1) were added, followed by vortexing for 20 s and centrifugation for 5 min at 1800g to achieve liquid-liquid extraction. The CHCl_3_ organic phase, containing the total lipid fraction, was gently retrieved and evaporated under a N_2_ stream. The dried lipophilic metabolites were then resuspended in 1 mL of BF_3_-MeOH, vortexed for 20 s, sonicated for 5 min with ice packs, and incubated for 10 min in a dry bath at 90 °C to allow derivatization. After this transesterification step, which makes the metabolites more volatile, 2 mL of CHCl_3_/H_2_O_Milli-Q_ (1:1) were added, samples were vortexed for 20 s and centrifuged for 5 min at 1800g. The CHCl_3_ phase was collected and transferred into injection vials for GC-MS analysis.

Aliquots of 5 µl were injected in splitless mode at 250°C in a gas chromatograph (7890B GC System-G3440B with the G4513A autosampler injector, Agilent Technologies) coupled to a mass selective detector (5977B MSD, Agilent Technologies). The transfer line temperature was set to 280°C. Metabolites were separated on an HP-5ms Ultra Inert column (30 m, 0.25 mm, and 0.25 μm, Agilent Technologies) with a helium mobile phase constant flow rate of 1 mL min^-1^. Metabolite mass spectra were acquired under electron ionization mode at 70 eV between 35 and 600 m/z set at a scan rate of 1.3 scan s^−1^. The GC run method consisted of three temperature ramps: starting at 60 °C, increasing by 25 °C min^-1^ up to 150 °C, then by 3 °C min^-1^ up to 230 °C, and finally by 30 °C min^-1^ up to 325 °C, followed by a 3-min post-run at 80 °C, for a total runtime of 36.4 min with the post-run.

A standard mix solution of C_8-C20_ and C_21-C40_ alkanes from Sigma Aldrich was injected with the same GC run method, to calculate the metabolite retention indexes (RI). Each experimental RI was calculated using the temperature programmed-Kovats index from Van Den Dool and Dec. Kratz, (1963), as follow:

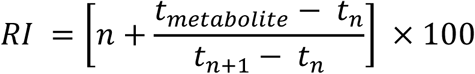

where *t_n_* and *t_n+1_* are the retention times of the reference *n-alkane* hydrocarbons eluting immediately before and after the target metabolite, and *n* is the carbon number of the trailing *n-alkane*.

Raw GC-MS data files (Agilent format) were converted to mzXML using ProteoWizard MSconvertGUI software (Chambers et al., 2012, v3.0.24100 64-bit), removing z-lib compression and vendor algorithm peak picking at MS level 1-2. The resulting outputs were pre-processed under R using the eRah package (Domingo-almenara et al., 2016, v2.0.1), with the following initial parameters: peak deconvolution (min.peak.width = 3, min.peak.height = 1500, noise.threshold = 1000, avoid. processing.mz = c(73, 149, 207)), peak alignment (min.spectra.cor = 0.75, max.time.dist = 5, mz. range = c(40:600), and missing compounds recovery (recMissComp, min.samples = 1). These settings optimized the balance between sensitivity and stringency, producing an initial metabolites dataset subsequently filtered, to remove peaks present in blanks (signal/noise ratio > 5), and metabolites detected in fewer than two samples. Duplicate or partially integrated peaks were manually verified and removed.

Metabolite annotation was performed using the NIST 2017 MS Search (v2.3) software coupled with InChI Library (v1.05), giving priority to the R_match_ NIST score and the match between the experimental RI and literature values. For each identified lipid metabolite, the relative proportion (as a percentage of the total lipid pool, %) was calculated from the integrated peak areas on the chromatograms. Metabolites subjected to BF_3_-MeOH transesterification and forming terminal methyl esters (-COOCH_3_), instead of their original free carboxylic acid group (such as fatty acid methyl esters, FAMEs), were corrected to represent their native pre-methylation molecular structures for more accurate biological interpretation. However, the reported R_match_ and RI values remain those of the methylated derivatives actually detected by GC-MS.

### Data treatment

Pigment and lipid relative proportion matrices (Hellinger-transformed) were combined using multiple factor analysis (MFA). Prior to this MFA, some red light lipid samples were excluded from the dataset as a precaution, due to non-exploitable chromatograms. Treatment-induced differences in multivariate compositions were then assessed across all MFA dimensions using permutational multivariate analysis of variance (PERMANOVA) based on Euclidean distances. Euclidean distances between treatment-defined group centroids in the MFA space were calculated across all dimensions and visualised using a heatmap to highlight multivariate dissimilarities. To assess treatment-induced structuring independently within pigment and lipid datasets, Hellinger-transformed matrices were analysed separately using principal component analysis (PCA), and Euclidean distances between treatment-defined centroids were computed across all PCA dimensions to quantify multivariate dissimilarities among treatments. The top 50% of metabolites most strongly correlated with, at least, one of the first two MFA axes (cos^2^ > 0.27) were selected for targeted, individual-level analysis to assess the effect of light treatments on their relative proportions using the Van der Waerden test. Based on Van der Waerden test results, metabolites were classified into four response clusters according to their variation across light treatments and time: Cluster 1, light-responsive metabolites without initial differences in dark-adapted controls; Cluster 2, metabolites showing both temporal and light-driven variation; Cluster 3, metabolites affected by temporal variation only; and Cluster 4, Non-responsive (stable) metabolites with no significant changes across time and treatments. Metabolites were then grouped by subclass using the ClassyFire classification and visualised in barplots to illustrate their distribution across clusters. Finally, treatment-induced changes in photosynthetic parameters were assessed between light treatments and their corresponding dark-adapted controls using Welch’s two-sample t-test.

## Results

### Light-driven and temporal remodeling of pigments and lipids in epipelic diatoms

The suspension derived from the natural winter microphytobenthic community, reached 4.5 ×10^6^ cells L^-1^ and was largely dominated by an assemblage of motile *Bacillariophyceae* with *Pleurosigma strigosum* accounted for 63.3%, followed by *Gyrosigma* sp. (13.3%), *Pleurosigma estuarii* (10%), and *Navicula* sp. (10%), while other pennate diatoms (e.g., *Entomoneis* sp. and *Cocconeis* sp.) contributed marginally (3.3%). Occasional nematodes and arthropods (e.g., copepods and nauplius larvae) were also observed.

Changes in diatom lipophilic metabolites were investigated through both lipophilic pigments and lipid composition. A total of 133 metabolites were identified, including 35 pigments and 98 lipids (Supplementary Table 2). Lipophilic pigments were dominated by chlorophyll *a* and its derivatives, fucoxanthin, and chlorophyll *c2*, accounting for 61.5%, 24.1%, and 5.9% of the average relative abundance, respectively. The remaining composition included unknown carotenoids, the diadinoxanthin + diatoxanthin pool, β-carotenes, and pheopigments (4.4%, 2.1%, 1.2%, and 0.7%, respectively; Supplementary Fig. 2A). Lipid composition was largely dominated by fatty acids (84.9%), with minor contributions including terpenes, alkanes, alkynes, and amino acids (6.7%, 4.7%, 1.9%, and 0.3%, respectively). At the individual metabolite level, total lipid composition was dominated by saturated fatty acids (14:0, 15:0, 16:0, and 18:0), which together accounted for more than 45.3%, followed by major polyunsaturated fatty acids (C16:3 ω4, C18:2 ω6, C18:3 ω6, and C20:5 ω3; 12.5%) and major monounsaturated fatty acids (C16:1 ω7, C16:1 ω5, and C17:1 ω7; 9%) (Supplementary Fig. 2B).

Based on their lipid and pigment compositions, samples show a significant structuring of their positions in multivariate space according to treatments (Fig. 1A and C; PERMANOVA on all MFA dimensions, R^2^ = 0.404, *p* = 0.001). The representation of the pigment and lipid groups along the first two MFA dimensions (Supplementary Fig. 3) revealed a shared structure of variation between the two data blocks. Dimension 1, associated mainly with differences in light spectra and temporal metabolic remodeling, showed a slightly stronger association with lipids, whereas Dimension 2, associated with differences in light intensity, was more strongly driven by pigment variation. Euclidean distances between sample group centroids across all MFA dimensions revealed a clear structure of pairwise dissimilarities among treatments, with red light treatments showing the strongest divergence from their corresponding dark-adapted controls than blue light treatments (Fig. 1B). This finding was confirmed when pigment and lipid datasets were analyzed separately, as PCA-based Euclidean distances revealed consistent structuring of light treatments. Lipids exhibited stronger differentiation under low light, particularly under red light compared to blue light, whereas pigments showed a similar red light effect combined with an increasing differentiation along the light intensity gradient (Supplementary Table 3). Nevertheless, in both lipid and pigment datasets, the contrast between low and high light intensities was more pronounced under blue light than under red light.

**Figure 1.**
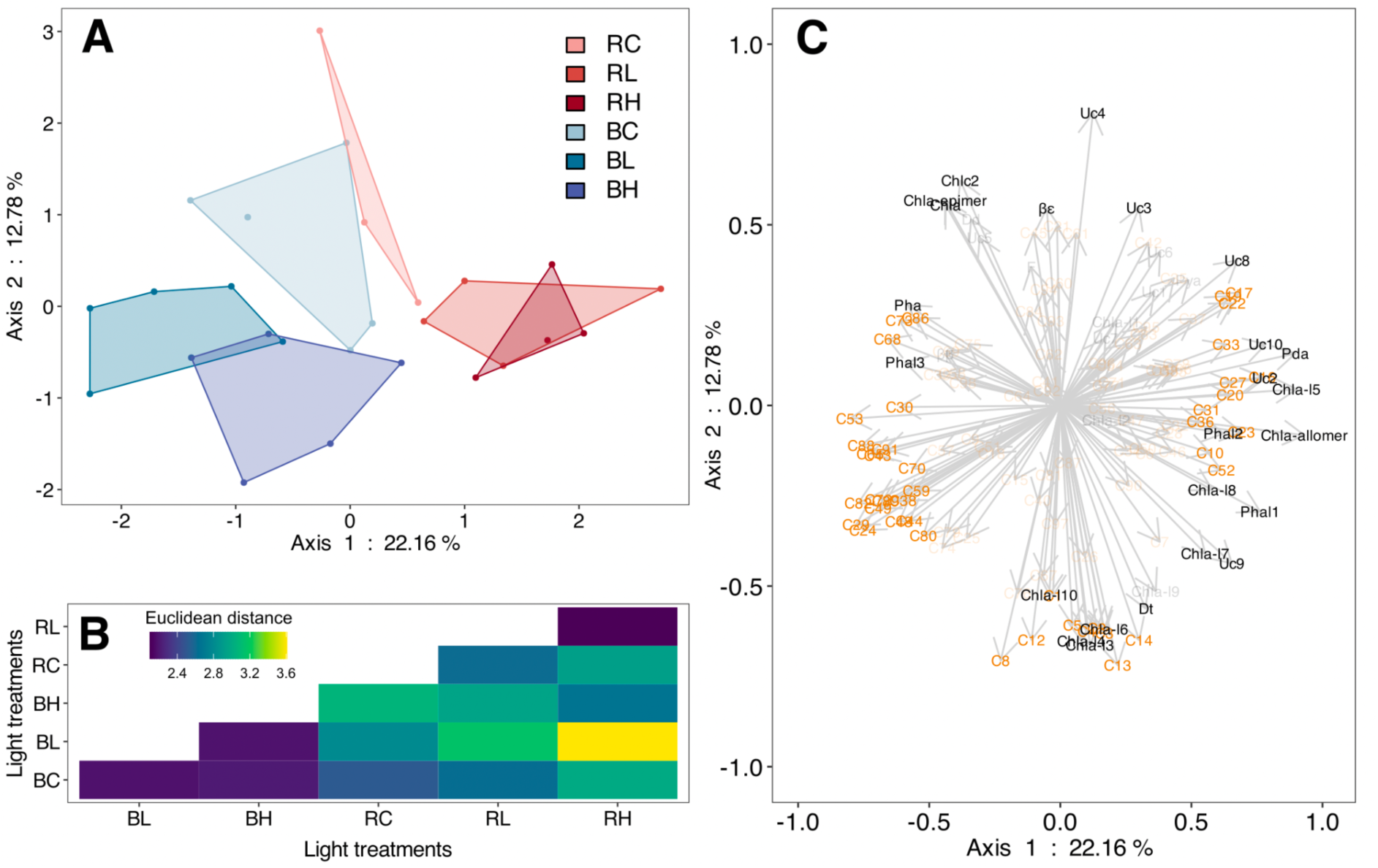
Untargeted lipid and lipophilic pigment responses to light treatments. **A.** Projection of samples in the factorial space derived from a Multiple Factor Analysis (MFA) based on Hellinger-transformed lipid and pigment composition (%) across treatments; **B.** Heatmap of Euclidean distances between group centroids performed across all MFA dimensions; **C.** Projection of variable vectors in the MFA factorial space, where their directions and lengths indicate their correlations and relative contributions to the structuring of the first two MFA axes (pigments and lipids abbreviations in Table s2). Only the top 50% (cos^2^ > 0.27) of metabolites most strongly correlated with, at least, one of the first two MFA axes being displayed for clarity. Light treatments: “Red” dark-adapted control, RC; Red low light, RL; Red high light, RH; “Blue” dark-adapted control, BC; Blue low light, BL; Blue high light, BH; (RC, n = 3; RL, RH, n = 4; BC, BL, BH, n = 5).

The top 50% of metabolites most strongly correlated with, at least, one of the first two MFA axes (cos^2^ > 0.27; *n* = 66 metabolites) were classified into four clusters according to their responses to light treatment and temporal metabolic variations (Fig. 2). Cluster 1 (*n* = 23; increasing to *n* = 38 when 100% of the identified metabolites were considered) comprised metabolites showing no differences between red and blue dark-adapted controls and exhibiting a significant response to at least one of the light treatments. This cluster notably revealed a decrease in the relative abundance of several polyunsaturated fatty acids along the red light intensity gradient, in contrast to an increase under blue low light (Fig. 3A). In particular, C16:4 exhibited a significant dynamic (Van der Waerden test, *p* < 0.05), as did C18:2 ω6, C18:4 ω3, and C20:4 ω3 (Supplementary Fig. 4A), the latter three belonging to the linoleic acid and derivative subclass, which was exclusively represented in this cluster (Fig. 2). Clusters 2 and 3 comprised metabolites showing temporal variation between red and blue dark-adapted controls, with or without a significant response under light treatments, but remained marginal containing only 4 and 8 metabolites (*n* = 16 and 8 when 100% of the identified metabolites were considered), respectively (Supplementary Fig. 4B and C). Finally, cluster 4 comprised the largest number of metabolites (*n* = 31; increasing to *n* = 71 when 100% of the identified metabolites were considered), characterized by overall stability throughout the experiment, with only weak emerging trends that were insufficient to support classification into the other three clusters (Supplementary Fig. 4D).

**Figure 2.**
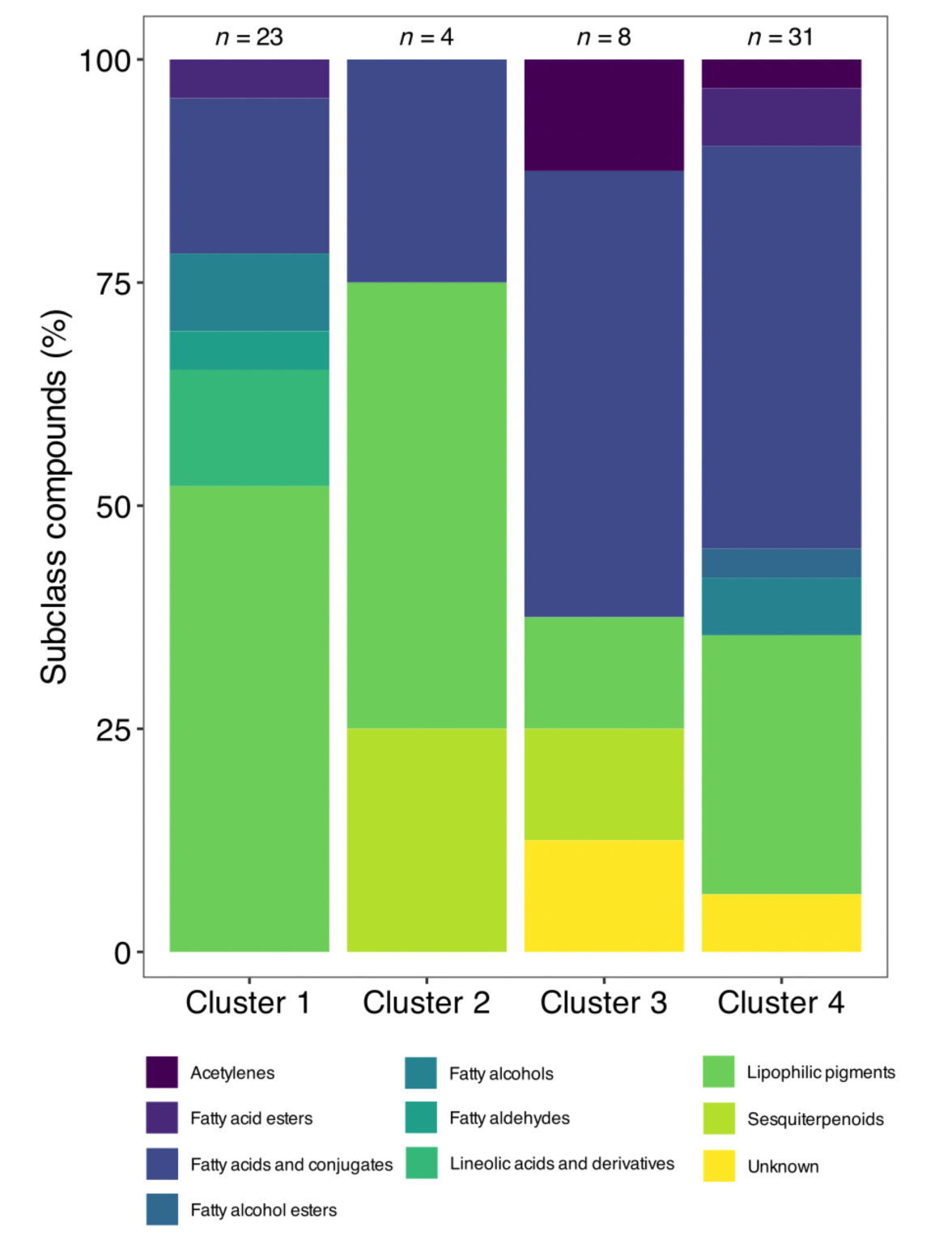
Metabolites subclass clusters according to temporal metabolic adjustments and light treatments. Barplots showing the relative proportions (%) of metabolite subclasses among the top 50% of metabolites most strongly correlated with, at least, one of the first two MFA axes (cos² > 0.27; n = 66). Metabolites were grouped into four clusters based on Van der Waerden test results, reflecting their response to light treatments and/or temporal variation: Cluster 1, light-responsive metabolites with no initial differences between dark-adapted controls; Cluster 2, metabolites showing both temporal variation and light responses; Cluster 3, metabolites driven by temporal variation only; and Cluster 4, stable metabolites with no significant changes across time and treatments.

**Figure 3.**
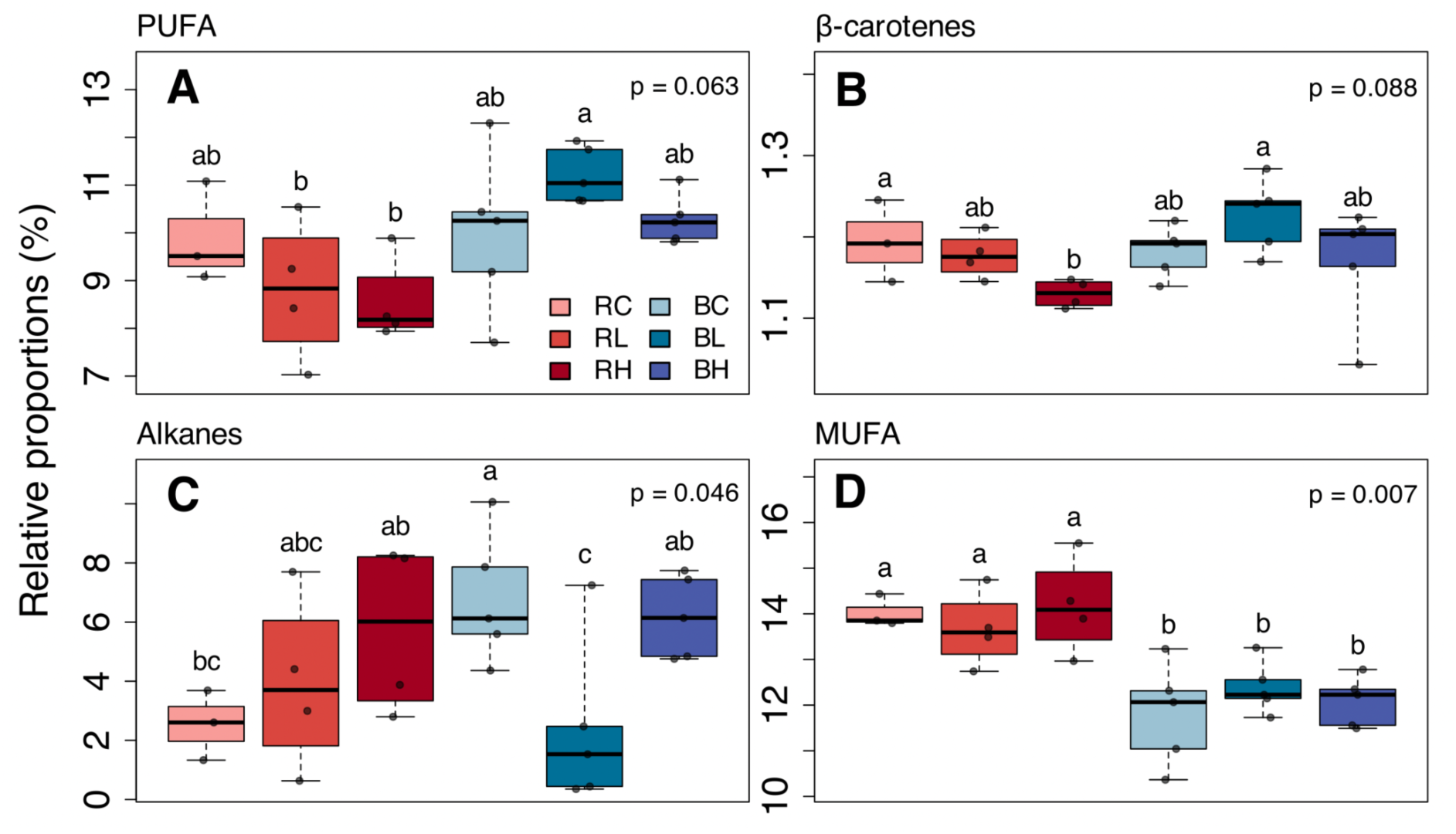
Untargeted key class lipid and lipophilic pigment responses to light treatments Boxplots showing the relative proportions (%) of the most relevant metabolites driving diatom metabolic responses under different light treatments, for **A.** Polyunsaturated fatty acids PUFAs from the top 50% of metabolites most strongly correlated with, at least, one of the first two MFA axes (cos² > 0.27); **B.** Total pool of β-carotenes; **C.** Total pool of alkanes and; **D.** Total pool of monounsaturated fatty acids MUFAs. Light treatments: “Red” dark-adapted control, RC; Red low light, RL; Red high light, RH; “Blue” dark-adapted control, BC; Blue low light, BL; Blue high light, BH. Groups formed according to light treatments were statistically tested using the Van der Waerden test (RC, n = 3; RL, RH, n = 4; BC, BL, BH, n = 5).

Examining the total pigment and lipid metabolite pools revealed class-level trends that were not apparent from individual metabolite dynamics. β-carotenes displayed response dynamics similar to those of polyunsaturated fatty acids under light treatments (Fig. 3B), primarily driven by the dominant β-β-carotene. In contrast, alkanes, the third most abundant lipid class, exhibited an exact opposite trend to polyunsaturated fatty acids and β-carotenes, with an increase along the red light intensity gradient and a significant decrease under blue low light (Van der Waerden test, *p* = 0.046; Fig. 3C), mainly driven by eicosane, hexacosane, docosane, and pentacosane. Finally, monounsaturated fatty acids showed no significant response to light treatments but were significantly influenced by temporal metabolic variation, as indicated by differences among controls (Van der Waerden test, *p* = 0.007; Fig. 3D).

### Xanthophyll-cycle and photophysiological parameters in epipelic diatoms

Within the overall pigment metabolic response, the xanthophylls diadinoxanthin and diatoxanthin, involved in the xanthophyll-cycle, were among the most responsive to light treatments (Fig. 4). Red and blue dark-adapted controls show a slight difference in diatoxanthin concentrations, resulting in a statistically different but comparable de-epoxidation state between the two treatments, with values of 3% and 2.6%, respectively (Supplementary Table 4). Diatoxanthin and diadinoxanthin concentrations displayed opposite responses under both light spectra, with diatoxanthin increasing and diadinoxanthin decreasing significantly along the intensity gradient (Van der Waerden test, *p* < 0.05; Fig. 4A). Their combined pool slightly declined with increasing light intensity except under red low light. Accordingly, the de-epoxidation state increased with light intensity under both spectra, with consistently higher values under red light, although remaining comparable to those under blue light, reaching maxima of 14.9% and 13.1%, respectively (Fig. 4B).

**Figure 4.**
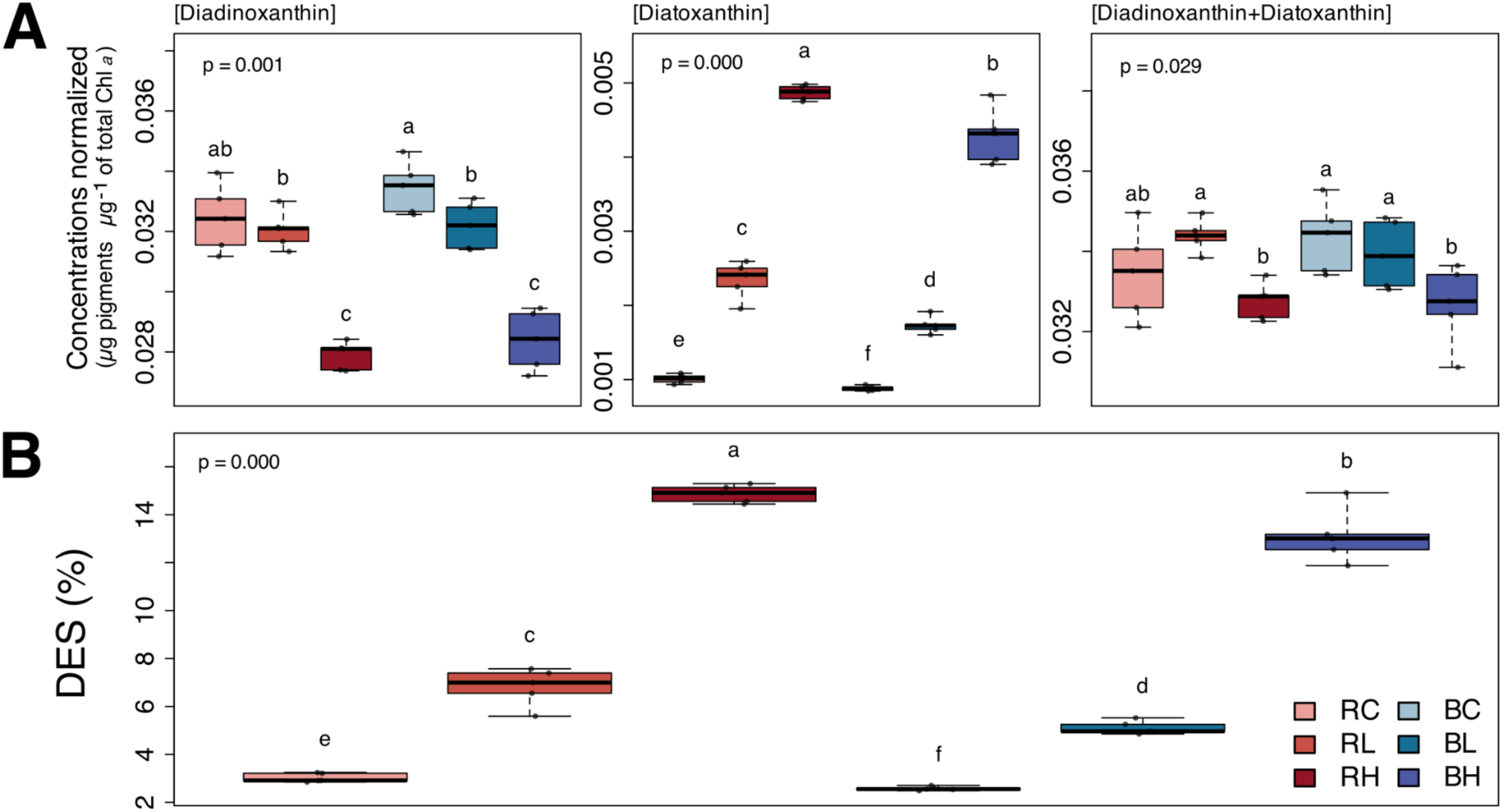
Xanthophyll-cycle responses of diatoms under different light treatments. **A.** Boxplots showing xanthophyll concentrations (*μ*g pigments *μ*g^-1^ Σ chlorophyll *a*) and; **B.** the de-epoxidation state (DES, %) under different light treatments. Differences between treatments were tested using the Van der Waerden test (n = 5). Light treatments: “Red” dark-adapted control, RC; Red low light, RL; Red high light, RH; “Blue” dark-adapted control, BC; Blue low light, BL; Blue high light, BH.

Considering the estimated portion of incident light actually absorbed by the cells, derived from reconstructed absorbance spectra based on sample pigment composition (Qphar) and further corrected for the cellular pigment packaging effect (Qpharpackage-corrected), together with the resulting cumulative absorbed photons (P_abs_) and absorbed energy (E_abs_) (Supplementary Table 5), de-epoxidation state responses exhibited markedly higher slopes under red than blue light. This could highlight a contrasting apparent xanthophyll-cycle response depending on the spectral light cue (Supplementary Fig. 5). The Qpharpackage-corrected values revealed that diatoms absorbed ∼4.2-fold more blue than red light. Consequently, the blue high light treatment corresponded to an energy input 3.9-fold higher than blue low light, 6.1-fold higher than red high light, and 22.3-fold higher than red low light.

In contrast to temporal metabolic changes, photosynthetic parameters remained stable, with no major temporal variations between red and blue dark-adapted samples, except for F_0_ (Supplementary Fig. 6). Dark-adapted cells exhibited initial Fv/Fm values ranging from 0.48 to 0.55 and α-slope values from 0.25 to 0.29 prior to light treatments (Supplementary Table 6).

Photophysiological changes clearly separated the blue high light treatment from the others with significant changes in parameters, including a +43.0% increase in Y(NO)_m obs_ and concomitant decreases in α-slope (−47.0%), Fv/Fm (−50.3%), and Y(NPQ)^1000^ (−42.5%) (Fig. 5 and Supplementary Fig. 7). In contrast, the three other light treatments induced only limited changes, with a slight increase in Y(NO)_m obs_ under red high light, and an increase in rETR_m obs_ under blue low light (Welch’s two-sample t-test, *p* < 0.05).

**Figure 5.**
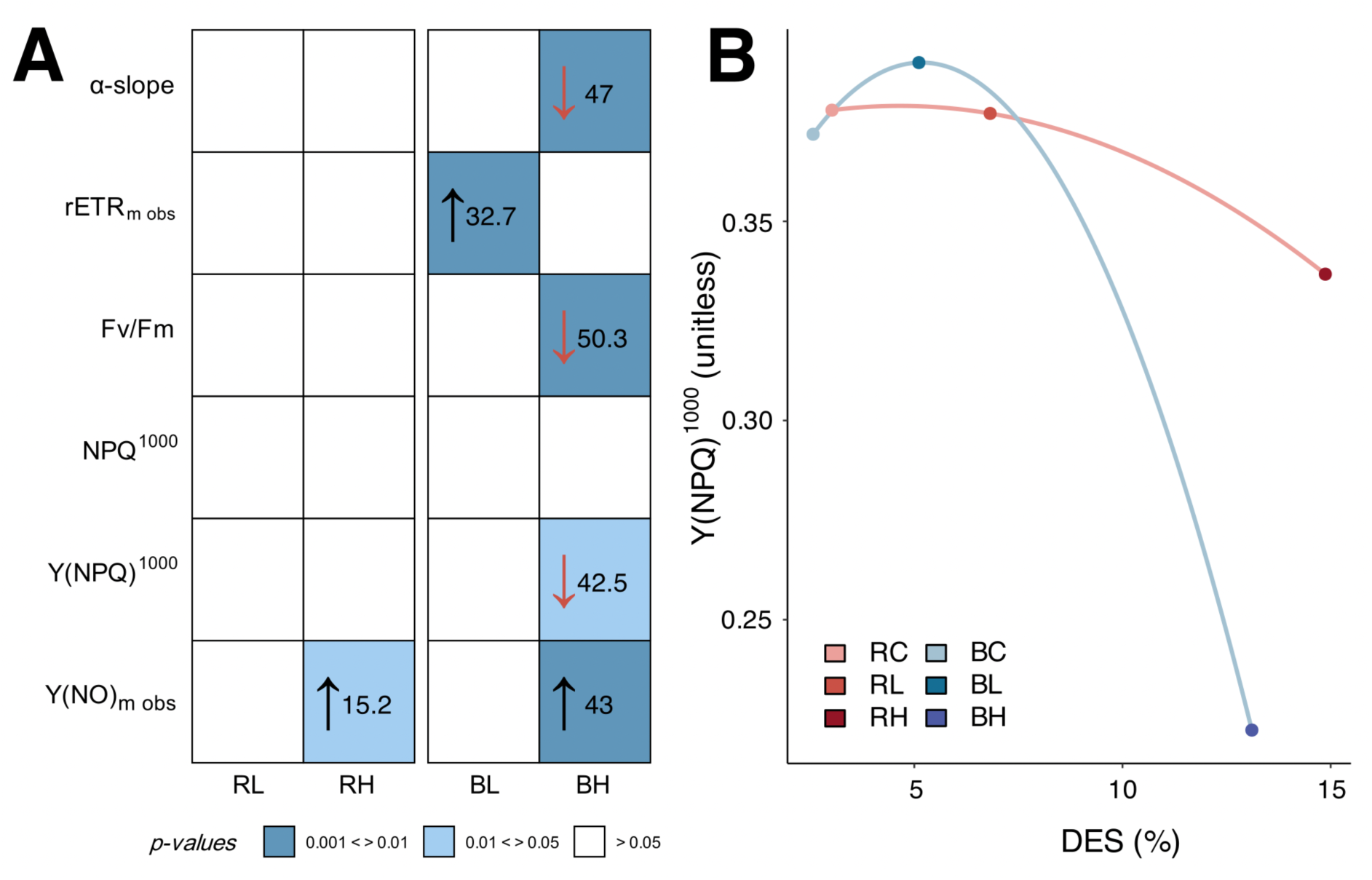
Cellular photosynthetic efficiency response across treatments. **A.** Heatmap of net photophysiological parameter responses under the light treatments. Displayed values (raw data presented in Supplementary Table 6), show the average change (%) between each light treatment and the red and blue dark-adapted controls, with arrows indicating the direction of change relative to the dark-adapted controls. Welch’s two-sample t-test (n = 3) was used to assess the treatment-induced changes ΔAfter treatment - Before treatment, relative to those observed in the dark-adapted controls. Alpha (arbitrary units) refers to the maximum light use coefficient for PSII at low light. Fv/Fm (unitless) corresponds to the maximum quantum yield of PSII in the dark-adapted state. rETRm obs (arbitrary units) and Y(NO)m obs (unitless) refer to the maximum values observed within the measured light curve range (without extrapolation). NPQ^1000^ (unitless) and Y(NPQ)^1000^ (unitless) correspond to the values observed at the highest light step of the light curve (1000 PPFD, without extrapolation); **B.** Plot showing the average photophysiological responses of Y(NPQ)^1000^ (n = 3) as a function of the average de-epoxidation states (%; n = 5). Light treatments: “Red” dark-adapted control, RC; Red low light, RL; Red high light, RH; “Blue” dark-adapted control, BC; Blue low light, BL; Blue high light, BH.

## Discussion

### Rapid lipophilic metabolite remodeling in epipelic diatoms

Lipophilic metabolite profiles revealed a clear chemotaxonomic signature characteristic of diatoms, marked by the enrichment of chlorophylls *a* and *c2*, fucoxanthin and xanthophylls, together with ω3- and ω6-polyunsaturated fatty acids, which are well-established pigment and lipid biomarkers of diatoms (Dunstan et al., 1993; Dijkman et al., 2010; Roy et al., 2011; Doose & Hubas, 2024). Combined with microscopic observations, these metabolic profiles corroborate an epipelic diatom community dominated by *Pleurosigma strigosum*, consistent with previous characterizations of muddy microphytobenthic assemblages (Underwood & Barnett, 2006; Méléder et al., 2007; Doose & Hubas, 2024). Given its dominant contribution to community biovolume (Desparmet et al., 2026), *P. strigosum* likely represents the main contributor to the metabolic and photosynthetic responses observed here.

Lipophilic metabolite profiling revealed contrasting metabolic remodeling driven by both temporal variation and light-dependent changes (Fig. 1–3). The temporal variation in monounsaturated fatty acids between red- and blue-light dark-adapted controls (Fig. 3D) illustrates this temporal component, which may reflect endogenous cellular regulation, as epipelic diatoms are known to tightly regulate multiple biological processes in anticipation of predictable changes, such as diel and tidal cycles (Consalvey et al., 2004; Coelho et al., 2011; Annunziata et al., 2019; Manzotti et al., 2025). Similarly, such endogenous metabolic adjustments may provide a fitness advantage by allowing diatoms to physiologically adjust to recurring environmental fluctuations, particularly by anticipating changes in light availability. However, temporal metabolic remodeling remained limited, with clusters 2 and 3 accounting for only ∼18% of the metabolites, compared with ∼81% in clusters 1 and 4, which were not significantly affected by temporal variation (Fig. 2).

Light treatments remained the main driver of the rapid lipophilic metabolite remodeling observed, with both light spectrum and intensity cues shaping metabolic trajectories and eliciting distinct metabolite-level responses. These results are consistent with previous studies showing that environmental changes can disrupt the balance between photosynthetic energy flux and carbon flux into biomass, leading to a reorganization of the partitioning of photosynthetically assimilated carbon among different macromolecular pools (e.g., proteins, lipids, carbohydrates, and nucleic acids) to sustain cellular function (Wagner et al., 2017; Doose and Hubas, 2024). In *Phaeodactylum tricornutum*, rapid shifts between red and blue light are known to trigger major changes in metabolome and macromolecular composition within minutes to hours (Jungandreas et al., 2014). Given these carbon partitioning strategies, the ability to rapidly adjust the pool of lipophilic metabolites within 30 min in response to different light cues is not surprising in epipelic diatoms inhabiting highly dynamic light environments (Kühl & Jørgensen, 1994; Jesus et al. 2009; Cartaxana et al., 2011; Underwood et al., 2022).

Among the key metabolites, polyunsaturated fatty acids and β-carotenes exhibited similar response dynamics across treatments, with contrasting changes reflecting distinct short-term photoacclimation strategies depending on the light cues received (Fig. 3A and B). Blue light is known to enhance carbon fixation and promote the synthesis of polyunsaturated fatty acid-enriched lipids, especially under low intensities that stimulate desaturation pathways (Guschina and Harwood, 2006; Chandrasekaran et al., 2014; Lima et al., 2021; Tanaka et al., 2022; Poliner et al., 2022; Svenning et al., 2024), consistent with the slight increase observed under blue low light (Fig. 3A). Fatty acids have been repeatedly identified in autotrophic organisms as highly responsive to various environmental stresses, including light, nutrients, temperature, pH, salinity, pollutants, metals, and CO_2_ availability (Los and Murata, 2004; Guschina and Harwood, 2006; Doose and Hubas 2024). Mainly found in membranes and lipid droplets, polyunsaturated fatty acids are essential metabolites involved in light-dependent growth strategies in diatoms (Zulu et al., 2018). Their rearrangement can lead to energy storage in triacylglycerols or to incorporation into structural lipids, where they contribute to thylakoid membrane integrity and fluidity, light-harvesting complex stability, and xanthophyll-cycle activity (Lepetit et al., 2010; Goss et al., 2020; Goss & Latowski, 2020; Büchel et al., 2022; Svenning et al., 2024). However, their high degree of unsaturation also makes them particularly susceptible to peroxidation under light stress (Doose et al., 2024), potentially constraining their accumulation under the blue high light treatment (Fig. 3A). The similar response dynamics of β-carotenes (Fig. 3B) likely reflect concurrent adjustments of the photosynthetic apparatus, as these accessory pigments play a dual role in optimizing light harvesting under favorable conditions and scavenging reactive oxygen species under light stress, while also serving as metabolic precursors for the biosynthesis of other pigments, including xanthophylls and fucoxanthin (Rijstenbil, 2003; Chandrasekaran et al., 2014; Kuczynska et al., 2015; Lima et al., 2021).

At the same time, alkanes exhibit the exact opposite response to polyunsaturated fatty acids and β-carotenes (Fig. 3C). These hydrocarbons are integral components of lipid metabolism, with biosynthetic pathways linked to those of fatty acids, occurring primarily through terminal enzymatic decarboxylation (C*_n_* to C*_n−1_*), which in some cases are directly regulated by blue light via fatty acid photodecarboxylase in microalgae (Herman and Zhang, 2016; Sorigué et al., 2017; Moulin et al., 2021). Even though *Bacillariophyceae* likely represent the major contribution to the metabolite pool, the origin of the alkanes observed here remains uncertain, as these metabolites are widely reported in both microalgae (Stonik and Stonik, 2015; López-Rosales et al., 2018; Harada et al., 2021; Manning, 2022) and associated phycosphere bacteria (Ladygina et al., 2006; Hagaggi & Abdul-Raouf, 2025), with such mixed microbial communities known to contribute to the observed chemical diversity (Hubas et al., 2018; Hubas et al., 2023; Helliwell et al., 2022). The alkane responses observed here may reflect either intrinsic alkane reorganization within diatoms as part of broader lipid remodeling, as well as a phycosphere bacterial response driven by changes in conditions and diatom activity (Gaubert-Boussarie et al., 2020; Doose and Hubas, 2024; Hubas et al., 2023; Helliwell et al., 2022). These saturated hydrocarbons, characterized by their chemical stability, may be associated with membrane structural functions, energy storage, growth and cell division, as well as cellular signalling (Ladygina et al., 2006; Lea-Smith et al., 2016; Sorigué et al., 2017; Manning, 2022; Doose and Hubas, 2024).

### Contrasting light spectra, comparable xanthophyll-cycle responses: different underlying mechanisms?

Despite the spectral- and intensity-dependent remodeling observed for lipophilic metabolites, xanthophyll-cycle responses appeared to be primarily structured by light intensity, with comparable de-epoxidation states under equivalent incident intensity regardless of light spectrum (Fig. 4). Although some studies have reported comparable de-epoxidation states in benthic diatoms under red and blue light exposure (McGee et al., 2020; Prins et al., 2020), with the total diadinoxanthin + diatoxanthin pool sometimes reaching higher values under red light conditions (Guérin et al., 2022), most studies conducted on pennate and centric diatoms have reported stronger de-epoxidation responses under blue light (Schellenberger Costa et al., 2013; Brunet et al., 2014; Valle et al., 2014; Smerilli et al., 2017; Su, 2019). The comparable de-epoxidation states observed here appear particularly unexpected when considering absorbed rather than incident light (Supplementary Fig. 5). Estimates based on Qphar, its package-effect correction, and the resulting cumulative absorbed photon flux (Pabs) and energy (Eabs) suggest that blue light was more efficiently absorbed and delivered a greater apparent energetic stimulus than red light (Supplementary Table 5). Blue light is known to be absorbed more efficiently, resulting in stronger photochemical excitation and greater energy dissipation requirements under high light (Brunet et al., 2014; Goessling et al., 2016), while also being intrinsically more photodamaging and associated with greater cellular oxidative stress (Goessling et al., 2016; Pashkovskiy et al., 2018). This stronger photophysiological impact of blue light appears to be consistent with the more pronounced alterations observed here under blue than red light (Fig. 5 and Supplementary Fig. 7).

Thus, comparable de-epoxidation states do not necessarily indicate equivalent physiological stimulation by the two light spectra, raising the question of why the xanthophyll-cycle response remained similar despite the apparently stronger stimulus under blue light. Moreover, the maximum de-epoxidation states observed here remained well below those commonly reported for epipelic diatoms (Van Leeuwe et al., 2008; Prins et al., 2020; Doose and Hubas, 2024), reaching up to 45% under full-spectrum light at 1500 µmol photons m^-2^ s^-1^ for 30 min (Desparmet et al., 2026).

Two non-mutually exclusive explanations may account for these de-epoxidation dynamics: an underperforming xanthophyll-cycle response under blue light and/or an enhanced response under red light.

### An enhanced xanthophyll-cycle response under red light?

One possibility is that red light could trigger regulatory mechanisms that enhance the photoprotective xanthophyll-cycle response, potentially through two non-mutually exclusive mechanisms. First, an imbalance in excitation between PSI and PSII, whereby red light may preferentially favor PSI excitation and promote cyclic electron flow around PSI, thereby increasing lumen acidification and consequently stimulating diadinoxanthin de-epoxidase activity (Munekage et al., 2004; Sato et al., 2014; Yamamoto and Shikanai, 2019). Second, red light photoreceptor signalling, including phytochrome-mediated regulation, may contribute to this enhanced response, given their increasingly recognized roles in diatom photoacclimation (Depauw et al., 2012; Lepetit and Dietzel, 2015; Fortunato et al., 2016; Jaubert et al., 2022; Duchêne et al., 2025; Font-Muñoz et al., 2026).

### An underperforming blue light response or a signature of karyostrophy?

A second possibility is that blue light induces an underperforming photoprotective xanthophyll-cycle response. One possible explanation is that monochromatic light may not fully engage the xanthophyll-cycle, which may require multiple wavelengths, as reported in *Pseudo-nitzschia multistriata* and *Skeletonema marinoi*, where photoprotection is enhanced by combining red and blue light (Brunet et al., 2014; Orefice et al., 2016). In *Skeletonema marinoi*, this partial response under monochromatic light is characterized by an initial concomitant increase in NPQ and de-epoxidation state under blue low light fluence, followed by a decline in NPQ and in the diadinoxanthin + diatoxanthin pool despite a higher de-epoxidation state under blue high light conditions (Chandrasekaran et al., 2014), consistent with our observations (Fig. 4 and 5B). In line with this and although blue light is known to effectively trigger photoacclimation mechanisms (Schellenberger Costa et al., 2013; Valle et al., 2014), the xanthophyll-cycle responses observed here under monochromatic blue light likely remain below the full capacity that can be engaged under a complete light spectrum, a conclusion that could also extend to monochromatic red light results.

A major additional factor to consider when interpreting these results is the ability of diatoms to modulate their functional absorption cross-section through karyostrophy and the resulting increase in pigment packaging, a mechanism particularly pronounced in *P. strigosum* (Furukawa et al., 1998; Bastos et al., 2025). Despite the corrections applied, Qphar estimates and the resulting Pabs and Eabs may not accurately capture the light actually experienced by the diatoms because karyostrophy is not accounted for. Consequently, estimated values (Supplementary Table 5), particularly under blue high light, are likely overestimated and should therefore be considered indicative rather than quantitative. Karyostrophy may also complicate the interpretation of fluorescence-based photophysiological measurements (Fig. 5), as chloroplast contraction alters chloroplast distribution and fluorescence signals without necessarily reflecting equivalent changes in photophysiological state (Bastos et al., 2025). Notably, the increase in Y(NO)_m obs_ under high intensity is therefore difficult to interpret, as it may result from karyostrophy, photoinhibition, or a combination of both (Nishiyama et al., 2006; Nishiyama et al., 2011; Waring et al., 2010; Foyer and Hanke, 2022). A similar interpretation may apply to the loss of Y(NPQ) under blue high light, which could reflect a mechanically reduced fluorescence signal associated with karyostrophy and/or a decoupling between diatoxanthin accumulation and NPQ, as large amounts of diatoxanthin may dissolve in the thylakoid membrane lipid phase, forming a lipid shield with antioxidant functions without contributing to quenching (Schumann et al., 2007; Van Leeuwe et al., 2008; Lepetit et al., 2010; Goss and Latowski, 2020; Croteau et al., 2021). Taken together, the xanthophyll-cycle response under blue light may not necessarily be underperforming, but could instead reflect strongly engaged karyostrophy, which may have reduced the effective light exposure experienced by the diatoms and thereby limited the need for a stronger de-epoxidation response.

Despite these considerations, an apparent paradox remains: blue low light showed stable Y(NO)_m obs_ and Y(NPQ)^1000^ parameters, whereas red high light showed increasing Y(NO)_m obs_ and decreasing Y(NPQ)^1000^, despite a 2.9-fold higher de-epoxidation state (Supplementary Table 4, 6 and Supplementary Fig. 7). If these responses result from differences in karyostrophy and/or photoinhibition, why would either process be more pronounced under red high light than under blue low light? In fact, one hypothesis to consider is a blue-light-dependent regulation of karyostrophy, as reported in centric diatoms by Furukawa et al. (1998). Here, blue low light may induce karyostrophy, potentially preventing photoinhibition and maintaining Y(NO)_m obs_ and Y(NPQ)^1000^, whereas karyostrophy may be weakly induced or absent under red high light, potentially resulting in photoinhibition. Evidence of such photoinhibition may lie in the observed decrease in β-carotenes (Fig. 3B), which may reflect their role in reactive oxygen species scavenging under red high light conditions. These mechanisms should also be considered in light of the observed lipophilic metabolite remodeling, particularly the increase in polyunsaturated fatty acids under blue low light, which may have contributed to maintaining photosynthetic performance (Fig. 3A).

### Lipophilic metabolite plasticity as a potential adaptative trait in diatoms facing fluctuating environmental niches

Beyond some general patterns, the responses observed here require careful consideration when interpreting their ecological significance, as diatom photoacclimation is highly species- and context-dependent, shaped by light history and environmental conditions (Waring et al., 2010; Mizrachi et al., 2019; McGee et al., 2020; Duarte et al., 2021; Jesus et al., 2023; Svenning et al., 2024). Moreover, our experimental setup imposed substantially different conditions on the diatoms from those experienced *in situ* (e.g., absence of sediment, resulting in reduced mobility, and monochromatic light). Nonetheless, the rapid lipophilic metabolite remodeling observed in these diatoms from a winter microphytobenthic community likely reflects, at least in part, intrinsic photoregulatory capacities shaped by its intertidal niche.

Diatoms can exploit spectral composition as an informative cue for photoregulation, with red-enriched spectra potentially indicating low tide and an increased likelihood of exposure to high and rapidly fluctuating light (Ragni, 2004; Waring et al., 2010; Brunet et al., 2014; Duchêne et al., 2025; Morelle et al., 2026). The stronger pigment and lipid remodeling observed under red light, together with the comparable xanthophyll-cycle response under red and blue light, may therefore suggest a particular responsivity to red wavelengths in these epipelic diatoms, potentially reflecting an adaptive strategy to anticipate changing light conditions. However, the plasticity observed under blue light, including the increase in polyunsaturated fatty acids under low light, further highlights the capacity to adjust metabolism according to the prevailing light environment, potentially reflecting trade-offs between photosynthetic performance, growth, and energy storage (Wagner et al., 2017; Croteau et al., 2021).

## Conclusion

The present work demonstrates that epipelic diatoms from a natural winter microphytobenthic community rapidly remodel their lipophilic metabolite composition, with contrasting responses depending on both spectral quality and light intensity. This substantial short-term metabolic plasticity suggests that adjustments in carbon allocation and photophysiological functioning according to the prevailing light environment may contribute to maintaining cellular homeostasis and photosynthetic performance. Specifically, red light induced the strongest remodelling of both pigment and lipid profiles, while eliciting a de-epoxidation state comparable to that observed under blue light at equivalent incident intensity, despite contrasting effects on apparent photosynthetic parameters. However, the cellular mechanisms underlying these responses remain unclear and require further investigation to determine their respective contributions and ecological significance. Overall, these findings highlight the capacity of epipelic diatoms to integrate spectral quality and light intensity cues through flexible metabolic and photophysiological responses, providing new insights into the diversity of photoregulatory strategies within the autotrophic microbiome. Such plasticity may contribute to the persistence of diatoms in highly variable environments such as intertidal mudflats.

## Author contributions

**AD** (Conceptualization, Methodology, Investigation, Data curation, Formal analysis, Visualization, Writing - original draft); **JL** (Formal analysis, Validation, Writing - review & editing); **BJ** (Data curation, Formal analysis, Validation, Writing - review & editing); **AM** (Investigation); and **CH** (Supervision, Methodology, Data curation, Formal analysis, Validation, Writing - review & editing).

## References

Annunziata, R., Ritter, A., Fortunato, A.E., Manzotti, A., Cheminant-Navarro, S., Agier, N., Huysman, M.J.J., Winge, P., Bones, A.M., Bouget, F.-Y., Cosentino Lagomarsino, M., Bouly, J.-P., Falciatore, A., 2019. bHLH-PAS protein RITMO1 regulates diel biological rhythms in the marine diatom Phaeodactylum tricornutum. Proc. Natl. Acad. Sci. U.S.A. 116, 13137–13142. 10.1073/pnas.1819660116

Bastos, A., Morelle, J., Frankenbach, S., Lavaud, J., Serôdio, J., 2025. Light response of karyostrophy in the benthic pennate diatom *Pleurosigma strigosum* (*Bacillariophyceae*): a complimentary photoprotective process? J Phycol 61, 1140– 1152. 10.1111/jpy.70055

Blommaert, L., 2017. Photoprotection in intertidal benthic diatoms. Ph.D. dissertation. Ghent University. Faculty of Sciences, Ghent, Belgium.

Blommaert, L., Huysman, M.J.J., Vyverman, W., Lavaud, J., Sabbe, K., 2017. Contrasting NPQ dynamics and xanthophyll cycling in a motile and a non-motile intertidal benthic diatom. Limnology & Oceanography 62, 1466–1479. 10.1002/lno.10511

Brotas, V., Plante-Cuny, M.-R., 2003. The use of HPLC pigment analysis to study microphytobenthos communities. Acta Oecol 24, S109–S115. 10.1016/S1146-609X(03)00013-4

Brunet, C., Chandrasekaran, R., Barra, L., Giovagnetti, V., Corato, F., Ruban, A.V., 2014. Spectral Radiation Dependent Photoprotective Mechanism in the Diatom *Pseudo-nitzschia multistriata*. PLoS ONE 9, e87015. 10.1371/journal.pone.0087015

Büchel, C., Goss, R., Bailleul, B., Campbell, D.A., Lavaud, J., Lepetit, B., 2022. Photosynthetic Light Reactions in Diatoms. I. The Lipids and Light-Harvesting Complexes of the Thylakoid Membrane, in: Falciatore, A., Mock, T. (Eds.), The Molecular Life of Diatoms. Springer International Publishing, Cham, pp. 397–422. 10.1007/978-3-030-92499-7_15

Cartaxana, P., Ruivo, M., Hubas, C., Davidson, I., Serôdio, J., Jesus, B., 2011. Physiological versus behavioral photoprotection in intertidal epipelic and epipsammic benthic diatom communities. J Exp Mar Biol Ecol 405, 120–127. 10.1016/j.jembe.2011.05.027

Chambers, M.C., Maclean, B., Burke, R., Amodei, D., Ruderman, D.L., Neumann, S., Gatto, L., Fischer, B., Pratt, B., Egertson, J., Hoff, K., Kessner, D., Tasman, N., Shulman, N., Frewen, B., Baker, T.A., Brusniak, M.-Y., Paulse, C., Creasy, D., Flashner, L., Kani, K., Moulding, C., Seymour, S.L., Nuwaysir, L.M., Lefebvre, B., Kuhlmann, F., Roark, J., Rainer, P., Detlev, S., Hemenway, T., Huhmer, A., Langridge, J., Connolly, B., Chadick, T., Holly, K., Eckels, J., Deutsch, E.W., Moritz, R.L., Katz, J.E., Agus, D.B., MacCoss, M., Tabb, D.L., Mallick, P., 2012. A cross-platform toolkit for mass spectrometry and proteomics. Nat Biotechnol 30, 918–920. 10.1038/nbt.2377

Chandrasekaran, R., Barra, L., Carillo, S., Caruso, T., Corsaro, M.M., Dal Piaz, F., Graziani, G., Corato, F., Pepe, D., Manfredonia, A., Orefice, I., Ruban, A.V., Brunet, C., 2014. Light modulation of biomass and macromolecular composition of the diatom Skeletonema marinoi. Journal of Biotechnology 192, 114–122. 10.1016/j.jbiotec.2014.10.016

Christof Klughammer, Ulrich Schreiber, 2008. Complementary PS II quantum yields calculated from simple fluorescence parameters measured by PAM fluorometry and the saturation pulse method. PAM Application Notes 1, 27–35.

Clementson, L.A., Wojtasiewicz, B., 2019. Dataset on the absorption characteristics of extracted phytoplankton pigments. Data Brief 24, 103875. 10.1016/j.dib.2019.103875

Coelho, H., Vieira, S., Serôdio, J., 2011. Endogenous versus environmental control of vertical migration by intertidal benthic microalgae. Eur J Phycol 46, 271–281. 10.1080/09670262.2011.598242

Consalvey, M., Paterson, D.M., Underwood, G.J.C., 2004. The ups and downs of life in a benthic biofilm: migration of benthic diatoms. Diatom Res 19, 181–202. 10.1080/0269249X.2004.9705870

Consalvey, M., Perkins, R.G., Paterson, D.M., Underwood, G.J.C., 2005. PAM fluorescence: a beginners guide for benthic diatoms. Diatom Res 20, 1–22. 10.1080/0269249X.2005.9705619

Croteau, D., Guérin, S., Bruyant, F., Ferland, J., Campbell, D.A., Babin, M., Lavaud, J., 2021. Contrasting nonphotochemical quenching patterns under high light and darkness aligns with light niche occupancy in Arctic diatoms. Limnology & Oceanography 66. 10.1002/lno.11587

Croteau, D., Lacour, T., Schiffrine, N., Morin, P., Forget, M., Bruyant, F., Ferland, J., Lafond, A., Campbell, D.A., Tremblay, J., Babin, M., Lavaud, J., 2022. Shifts in growth light optima among diatom species support their succession during the spring bloom in the Arctic. Journal of Ecology 110, 1356–1375. 10.1111/1365-2745.13874

Depauw, F.A., Rogato, A., Ribera d’Alcala, M., Falciatore, A., 2012. Exploring the molecular basis of responses to light in marine diatoms. Journal of Experimental Botany 63, 1575–1591. 10.1093/jxb/ers005

Desparmet, A., Jesus, B., Robinet, T., Dufour, T., Hubas, C., 2026. Reactive oxygen species trigger downward vertical migration in diatom microphytobenthic biofilms as a strategy to cope with oxidative stress. The ISME Journal 20, wrag034. 10.1093/ismejo/wrag034

Dijkman, N.A., Boschker, H.T.S., Stal, L.J., Kromkamp, J.C., 2010. Composition and heterogeneity of the microbial community in a coastal microbial mat as revealed by the analysis of pigments and phospholipid-derived fatty acids. Journal of Sea Research 63, 62–70. 10.1016/j.seares.2009.10.002

Domingo-Almenara, X., Brezmes, J., Vinaixa, M., Samino, S., Ramirez, N., Ramon-Krauel, M., Lerin, C., Díaz, M., Ibáñez, L., Correig, X., Perera-Lluna, A., Yanes, O., 2016. eRah: A Computational Tool Integrating Spectral Deconvolution and Alignment with Quantification and Identification of Metabolites in GC/MS-Based Metabolomics. Anal. Chem. 88, 9821–9829. 10.1021/acs.analchem.6b02927

Doose, C., Hubas, C., 2024. The metabolites of light: Untargeted metabolomic approaches bring new clues to understand light-driven acclimation of intertidal mudflat biofilm. Science of The Total Environment 912, 168692. 10.1016/j.scitotenv.2023.168692

Doose, C., Oger, C., Mas-Normand, L., Durand, T., Hubas, C., 2024. Non-enzymatic oxylipin production in a mudflat microphytobenthic biofilm: evidence of a diatom response to light. Front. Photobiol. 2, 1441713. 10.3389/fphbi.2024.1441713

Duarte, B., Feijão, E., Goessling, J.W., Caçador, I., Matos, A.R., 2021. Pigment and Fatty Acid Production under Different Light Qualities in the Diatom *Phaeodactylum tricornutum*. Applied Sciences 11, 2550. 10.3390/app11062550

Duchêne, C., Bouly, J.-P., Pierella Karlusich, J.J., Vernay, E., Sellés, J., Bailleul, B., Bowler, C., Ribera d’Alcalà, M., Falciatore, A., Jaubert, M., 2025. Diatom phytochromes integrate the underwater light spectrum to sense depth. Nature 637, 691–697. 10.1038/s41586-024-08301-3

Dunstan, G.A., Volkman, J.K., Barrett, S.M., Leroi, J.-M., Jeffrey, S.W., 1993. Essential polyunsaturated fatty acids from 14 species of diatom (*Bacillariophyceae*). Phytochemistry 35, 155–161. 10.1016/S0031-9422(00)90525-9

Eaton, J.W., Moss, B., 1966. The estimation of numbers and pigment content in epipelic algal populations. Limnology & Oceanography 11, 584–595. 10.4319/lo.1966.11.4.0584

Eilers, P.H.C., Peeters, J.C.H., 1988. A model for the relationship between light intensity and the rate of photosynthesis in phytoplankton. Ecol Modell 42, 199–215. 10.1016/0304-3800(88)90057-9

Falciatore, A., d’Alcalà, M.R., Croot, P., Bowler, C., 2000. Perception of Environmental Signals by a Marine Diatom. Science 288, 2363–2366. 10.1126/science.288.5475.2363

Falciatore, A., Jaubert, M., Bouly, J.-P., Bailleul, B., Mock, T., 2020. Diatom Molecular Research Comes of Age: Model Species for Studying Phytoplankton Biology and Diversity. Plant Cell 32, 547–572. 10.1105/tpc.19.00158

Finkel, Z.V., Follows, M.J., Liefer, J.D., Brown, C.M., Benner, I., Irwin, A.J., 2016. Phylogenetic Diversity in the Macromolecular Composition of Microalgae. PLoS ONE 11, e0155977. 10.1371/journal.pone.0155977

Font-Muñoz, J.S., Jaubert, M., Sourisseau, M., Tuval, I., Bailleul, B., Duchêne, C., Basterretxea, G., Falciatore, A., 2026. Phytochromes facilitate social behaviour in marine diatoms. Nat Commun. 10.1038/s41467-026-70219-3

Fortunato, A.E., Jaubert, M., Enomoto, G., Bouly, J.-P., Raniello, R., Thaler, M., Malviya, S., Bernardes, J.S., Rappaport, F., Gentili, B., Huysman, M.J.J., Carbone, A., Bowler, C., d’Alcalà, M.R., Ikeuchi, M., Falciatore, A., 2016. Diatom Phytochromes Reveal the Existence of Far-Red-Light-Based Sensing in the Ocean. Plant Cell 28, 616–628. 10.1105/tpc.15.00928

Foyer, C.H., Hanke, G., 2022. ROS production and signalling in chloroplasts: cornerstones and evolving concepts. Plant J 111, 642–661. 10.1111/tpj.15856

Furukawa, T., Watanabe, M., Shihira-Ishikawa, I., 1998. Green- and blue-light-mediated chloroplast migration in the centric diatom *Pleurosira laevis*. Protoplasma 203, 214–220. 10.1007/BF01279479

Gaubert-Boussarie, J., Prado, S., Hubas, C., 2020. An untargeted metabolomic approach for microphytobenthic biofilms in intertidal mudflats. Front. Mar. Sci. 7, 250. 10.3389/fmars.2020.00250

Gilbert, M., Domin, A., Becker, A., Wilhelm, C., 2000. Estimation of Primary Productivity by Chlorophyll a in vivo Fluorescence in Freshwater Phytoplankton. Photosynt. 38, 111–126. 10.1023/A:1026708327185

Goessling, J.W., Cartaxana, P., Kühl, M., 2016. Photo-Protection in the Centric Diatom Coscinodiscus granii is Not Controlled by Chloroplast High-Light Avoidance Movement. Front. Mar. Sci. 2. 10.3389/fmars.2015.00115

Goss, R., Latowski, D., 2020. Lipid Dependence of Xanthophyll Cycling in Higher Plants and Algae. Front. Plant Sci. 11, 455. 10.3389/fpls.2020.00455

Guérin, S., Raguénès, L., Croteau, D., Babin, M., Lavaud, J., 2022. Potential for the Production of Carotenoids of Interest in the Polar Diatom Fragilariopsis cylindrus. Marine Drugs 20, 491. 10.3390/md20080491

Guschina, I.A., Harwood, J.L., 2006. Lipids and lipid metabolism in eukaryotic algae. Progress in Lipid Research 45, 160–186. 10.1016/j.plipres.2006.01.001

Hagaggi, N.Sh.A., Abdul-Raouf, U.M., 2025. Recent trends in microbial production of alkanes. World J Microbiol Biotechnol 41, 320. 10.1007/s11274-025-04536-y

Harada, N., Hirose, Y., Chihong, S., Kurita, H., Sato, M., Onodera, J., Murata, K., Itoh, F., 2021. A novel characteristic of a phytoplankton as a potential source of straight-chain alkanes. Sci Rep 11, 14190. 10.1038/s41598-021-93204-w

Helliwell, K.E., Shibl, A.A., Amin, S.A., 2022. The Diatom Microbiome: New Perspectives for Diatom-Bacteria Symbioses, in: Falciatore, A., Mock, T. (Eds.), The Molecular Life of Diatoms. Springer International Publishing, Cham, pp. 679–712. 10.1007/978-3-030-92499-7_23

Herman, N.A., Zhang, W., 2016. Enzymes for fatty acid-based hydrocarbon biosynthesis. Current Opinion in Chemical Biology 35, 22–28. 10.1016/j.cbpa.2016.08.009

Hubas, C., Gaubert-Boussarie, J., D’Hondt, A.-S., Jesus, B., Lamy, D., Meleder, V., Prins, A., Rosa, P., Stock, W., Sabbe, K., 2023. Identification of microbial exopolymer producers in sandy and muddy intertidal sediments by compound-specific isotope analysis. Peer Community Journal 3, e104. 10.24072/pcjournal.336

Hubas, C., Passarelli, C., Paterson, D.M., 2018. Microphytobenthic biofilms: composition and interactions, in: Beninger, P.G. (Ed.), Mudflat Ecology. Springer International Publishing, Cham, pp. 63–90. 10.1007/978-3-319-99194-8_4

Jaubert, M., Duchêne, C., Kroth, P.G., Rogato, A., Bouly, J.-P., Falciatore, A., 2022. Sensing and Signalling in Diatom Responses to Abiotic Cues, in: Falciatore, A., Mock, T. (Eds.), The Molecular Life of Diatoms. Springer International Publishing, Cham, pp. 607–639. 10.1007/978-3-030-92499-7_21

Jesus, B., Brotas, V., Ribeiro, L., Mendes, C.R., Cartaxana, P., Paterson, D.M., 2009. Adaptations of microphytobenthos assemblages to sediment type and tidal position. Cont Shelf Res 29, 1624–1634. 10.1016/j.csr.2009.05.006

Jesus, B., Jauffrais, T., Trampe, E., Méléder, V., Ribeiro, L., Bernhard, J.M., Geslin, E., Kühl, M., 2023. Microscale imaging sheds light on species-specific strategies for photo-regulation and photo-acclimation of microphytobenthic diatoms. Environ Microbiol 25, 3087–3103. 10.1111/1462-2920.16499

Jungandreas, A., Schellenberger Costa, B., Jakob, T., Von Bergen, M., Baumann, S., Wilhelm, C., 2014. The Acclimation of *Phaeodactylum tricornutum* to Blue and Red Light Does Not Influence the Photosynthetic Light Reaction but Strongly Disturbs the Carbon Allocation Pattern. PLoS ONE 9, e99727. 10.1371/journal.pone.0099727

Kuczynska, P., Jemiola-Rzeminska, M., Strzalka, K., 2015. Photosynthetic pigments in diatoms. Mar Drugs 13, 5847–5881. 10.3390/md13095847

Kühl, M., Jorgensen, B.B., 1994. The light field of microbenthic communities: radiance distribution and microscale optics of sandy coastal sediments. Limnol. Oceanogr. 39, 1368–1398. 10.4319/lo.1994.39.6.1368

Ladygina, N., Dedyukhina, E.G., Vainshtein, M.B., 2006. A review on microbial synthesis of hydrocarbons. Process Biochemistry 41, 1001–1014. 10.1016/j.procbio.2005.12.007

Laviale, M., Frankenbach, S., Serôdio, J., 2016. The importance of being fast: comparative kinetics of vertical migration and non-photochemical quenching of benthic diatoms under light stress. Mar Biol 163, 10. 10.1007/s00227-015-2793-7

Lea-Smith, D.J., Ortiz-Suarez, M.L., Lenn, T., Nürnberg, D.J., Baers, L.L., Davey, M.P., Parolini, L., Huber, R.G., Cotton, C.A.R., Mastroianni, G., Bombelli, P., Ungerer, P., Stevens, T.J., Smith, A.G., Bond, P.J., Mullineaux, C.W., Howe, C.J., 2016. Hydrocarbons Are Essential for Optimal Cell Size, Division, and Growth of Cyanobacteria. Plant Physiol. 172, 1928–1940. 10.1104/pp.16.01205

Lepetit, B., Campbell, D.A., Lavaud, J., Büchel, C., Goss, R., Bailleul, B., 2022. Photosynthetic Light Reactions in Diatoms. II. The Dynamic Regulation of the Various Light Reactions, in: Falciatore, A., Mock, T. (Eds.), The Molecular Life of Diatoms. Springer International Publishing, Cham, pp. 423–464. 10.1007/978-3-030-92499-7_16

Lepetit, B., Dietzel, L., 2015. Light signaling in photosynthetic eukaryotes with ‘green’ and ‘red’ chloroplasts. Environ Exp Bot 114, 30–47. 10.1016/j.envexpbot.2014.07.007

Lepetit, B., Volke, D., Gilbert, M., Wilhelm, C., Goss, R., 2010. Evidence for the existence of one antenna-associated, lipid-dissolved and two protein-bound pools of diadinoxanthin cycle pigments in diatoms. Plant Physiol 154, 1905–1920. 10.1104/pp.110.166454

Lima, S., Schulze, P.S.C., Schüler, L.M., Rautenberger, R., Morales-Sánchez, D., Santos, T.F., Pereira, H., Varela, J.C.S., Scargiali, F., Wijffels, R.H., Kiron, V., 2021. Flashing light emitting diodes (LEDs) induce proteins, polyunsaturated fatty acids and pigments in three microalgae. Journal of Biotechnology 325, 15–24. 10.1016/j.jbiotec.2020.11.019

López-Rosales, A.R., Ancona-Canché, K., Chavarria-Hernandez, J.C., Barahona-Pérez, F., Toledano-Thompson, T., Garduño-Solórzano, G., López-Adrian, S., Canto-Canché, B., Polanco-Lugo, E., Valdez-Ojeda, R., 2018. Fatty Acids, Hydrocarbons and Terpenes of Nannochloropsis and Nannochloris Isolates with Potential for Biofuel Production. Energies 12, 130. 10.3390/en12010130

Los, D.A., Murata, N., 2004. Membrane fluidity and its roles in the perception of environmental signals. Biochimica et Biophysica Acta (BBA) - Biomembranes 1666, 142–157. 10.1016/j.bbamem.2004.08.002

Malerba, M.E., Palacios, M.M., Palacios Delgado, Y.M., Beardall, J., Marshall, D.J., 2018. Cell size, photosynthesis and the package effect: an artificial selection approach. New Phytologist 219, 449–461. 10.1111/nph.15163

Manning, S.R., 2022. Microalgal lipids: biochemistry and biotechnology. Current Opinion in Biotechnology 74, 1–7. 10.1016/j.copbio.2021.10.018

Manzotti, A., Monteil, R., Cheminant Navarro, S., Croteau, D., Charreton, L., Hoguin, A., Strumpen, N.F., Jallet, D., Daboussi, F., Kroth, P.G., Bouget, F., Jaubert, M., Bailleul, B., Bouly, J., Falciatore, A., 2025. Circadian regulation of key physiological processes by the RITMO1 clock protein in the marine diatom *Phaeodactylum tricornutum*. New Phytologist 246, 1724–1739. 10.1111/nph.70099

McGee, D., Archer, L., Fleming, G.T.A., Gillespie, E., Touzet, N., 2020. Influence of spectral intensity and quality of LED lighting on photoacclimation, carbon allocation and high-value pigments in microalgae. Photosynth Res 143, 67–80. 10.1007/s11120-019-00686-x

Méléder, V., Rincé, Y., Barillé, L., Gaudin, P., Rosa, P., 2007. Spatiotemporal changes in microphytobenthos assemblages in a macrotidal flat (Bourgneuf Bay,France). J Phycol 43, 1177–1190. 10.1111/j.1529-8817.2007.00423.x

Mizrachi, A., Graff Van Creveld, S., Shapiro, O.H., Rosenwasser, S., Vardi, A., 2019. Light-dependent single-cell heterogeneity in the chloroplast redox state regulates cell fate in a marine diatom. eLife 8, e47732. 10.7554/eLife.47732

Morelle, J., Bastos, A., Frankenbach, S., Frommlet, J.C., Campbell, D.A., Lavaud, J., Serôdio, J., 2024. The photoprotective behavior of a motile benthic diatom as elucidated from the interplay between cell motility and physiological responses to a light microgradient using a novel experimental setup. Microb Ecol 87, 40. 10.1007/s00248-024-02354-7

Morelle, J., Bastos, A., Pereira, L.F., Frankenbach, S., Lavaud, J., Serôdio, J., 2026. Far-red light regulates phototactic behavior of benthic pennate epipelic diatoms under low irradiance. Photosynth Res 164, 4. 10.1007/s11120-025-01195-w

Moulin, S.L.Y., Beyly-Adriano, A., Cuiné, S., Blangy, S., Légeret, B., Floriani, M., Burlacot, A., Sorigué, D., Samire, P.-P., Li-Beisson, Y., Peltier, G., Beisson, F., 2021. Fatty acid photodecarboxylase is an ancient photoenzyme that forms hydrocarbons in the thylakoids of algae. Plant Physiology 186, 1455–1472. 10.1093/plphys/kiab168

Munekage, Y., Hashimoto, M., Miyake, C., Tomizawa, K.-I., Endo, T., Tasaka, M., Shikanai, T., 2004. Cyclic electron flow around photosystem I is essential for photosynthesis. Nature 429, 579–582. 10.1038/nature02598

Nishiyama, Y., Allakhverdiev, S.I., Murata, N., 2011. Protein synthesis is the primary target of reactive oxygen species in the photoinhibition of photosystem II. Physiologia Plantarum 142, 35–46. 10.1111/j.1399-3054.2011.01457.x

Nishiyama, Y., Allakhverdiev, S.I., Murata, N., 2006. A new paradigm for the action of reactive oxygen species in the photoinhibition of photosystem II. Biochim Biophys Acta Bioenerg 1757, 742–749. 10.1016/j.bbabio.2006.05.013

Orefice, I., Chandrasekaran, R., Smerilli, A., Corato, F., Caruso, T., Casillo, A., Corsaro, M.M., Piaz, F.D., Ruban, A.V., Brunet, C., 2016. Light-induced changes in the photosynthetic physiology and biochemistry in the diatom Skeletonema marinoi. Algal Research 17, 1–13. 10.1016/j.algal.2016.04.013

Pashkovskiy, P.P., Soshinkova, T.N., Korolkova, D.V., Kartashov, A.V., Zlobin, I.E., Lyubimov, V.Yu., Kreslavski, V.D., Kuznetsov, Vl.V., 2018. The effect of light quality on the pro-/antioxidant balance, activity of photosystem II, and expression of light-dependent genes in Eutrema salsugineum callus cells. Photosynth Res 136, 199–214. 10.1007/s11120-017-0459-7

Passarelli, C., Meziane, T., Thiney, N., Boeuf, D., Jesus, B., Ruivo, M., Jeanthon, C., Hubas, C., 2015. Seasonal variations of the composition of microbial biofilms in sandy tidal flats: Focus of fatty acids, pigments and exopolymers. Estuarine, Coastal and Shelf Science 153, 29–37. 10.1016/j.ecss.2014.11.013

Perkins, R., Lavaud, J., Serôdio, J., Mouget, J., Cartaxana, P., Rosa, P., Barille, L., Brotas, V., Jesus, B., 2010. Vertical cell movement is a primary response of intertidal benthic biofilms to increasing light dose. Mar. Ecol. Prog. Ser. 416, 93–103. 10.3354/meps08787

Perkins, R., Underwood, G., Brotas, V., Snow, G., Jesus, B., Ribeiro, L., 2001. Responses of microphytobenthos to light: primary production and carbohydrate allocation over an emersion period. Mar. Ecol. Prog. Ser. 223, 101–112. 10.3354/meps223101

Pierella Karlusich, J.J., Cosnier, K., Zinger, L., Henry, N., Nef, C., Bernard, G., Scalco, E., Dvorak, E., Tara Oceans Coordinators, Acinas, S.G., Babin, M., Bork, P., Boss, E., Bowler, C., Cochrane, G., De Vargas, C., Gorsky, G., Grimsley, N., Guidi, L., Iudicone, D., Jaillon, O., Kandels, S., Karp-Boss, L., Karsenti, E., Not, F., Ogata, H., Pesant, S., Poulton, N., Sardet, C., Speich, S., Stemmann, L., Sullivan, M.B., Sunagawa, S., Wincker, P., Rocha Jimenez Vieira, F., Delage, E., Chaffron, S., Ovchinnikov, S., Zingone, A., Bowler, C., 2025. Patterns and drivers of diatom diversity and abundance in the global ocean. Nat Commun 16, 3452. 10.1038/s41467-025-58027-7

Poliner, E., Busch, A.W.U., Newton, L., Kim, Y.U., Clark, R., Gonzalez-Martinez, S.C., Jeong, B.-R., Montgomery, B.L., Farré, E.M., 2022. Aureochromes maintain polyunsaturated fatty acid content in *Nannochloropsis oceanica*. Plant Physiology 189, 906–921. 10.1093/plphys/kiac052

Prins, A., Deleris, P., Hubas, C., Jesus, B., 2020. Effect of light intensity and light quality on diatom behavioral and physiological photoprotection. Front. Mar. Sci. 7, 203. 10.3389/fmars.2020.00203

Ragni, R., Cicco, S., Vona, D., Leone, G., Farinola, G.M., 2017. Biosilica from diatoms microalgae: smart materials from bio-medicine to photonics. J. Mater. Res. 32, 279–291. 10.1557/jmr.2016.459

Rijstenbil, J., 2003. Effects of UVB radiation and salt stress on growth, pigments and antioxidative defence of the marine diatom Cylindrotheca closterium. Mar. Ecol. Prog. Ser. 254, 37–48. 10.3354/meps254037

Roy, S., Llewellyn, C.A., Egeland, E.S., Johnsen, G. (Eds.), 2011. Phytoplankton pigments: characterization, chemotaxonomy and applications in oceanography. Cambridge University Press. 10.1017/CBO9780511732263

Sato, R., Ohta, H., Masuda, S., 2014. Prediction of respective contribution of linear electron flow and PGR5-dependent cyclic electron flow to non-photochemical quenching induction. Plant Physiology and Biochemistry 81, 190–196. 10.1016/j.plaphy.2014.03.017

Schellenberger Costa, B., Jungandreas, A., Jakob, T., Weisheit, W., Mittag, M., Wilhelm, C., 2013. Blue light is essential for high light acclimation and photoprotection in the diatom *Phaeodactylum tricornutum*. Journal of Experimental Botany 64, 483–493. 10.1093/jxb/ers340

Schumann, A., Goss, R., Jakob, T., Wilhelm, C., 2007. Investigation of the quenching efficiency of diatoxanthin in cells of *Phaeodactylum tricornutum* (*Bacillariophyceae*) with different pool sizes of xanthophyll cycle pigments. Phycologia 46, 113–117. 10.2216/06-30.1

Serôdio, J., Ezequiel, Barnett, A., Mouget, Méléder, V., Laviale Lavaud, J., 2012. Efficiency of photoprotection in microphytobenthos: role of vertical migration and the xanthophyll cycle against photoinhibition. Aquat Microb Ecol 16, 161–175. 10.3354/ame01591

Shimakawa, G., Miyake, C., 2018. Oxidation of P700 Ensures Robust Photosynthesis. Front. Plant Sci. 9, 1617. 10.3389/fpls.2018.01617

Silsbe, G.M., Kromkamp, J.C., 2012. Modeling the irradiance dependency of the quantum efficiency of photosynthesis. Limnol Oceanogr Methods 10, 645–652. 10.4319/lom.2012.10.645

Smerilli, A., Orefice, I., Corato, F., Gavalás Olea, A., Ruban, A.V., Brunet, C., 2017. Photoprotective and antioxidant responses to light spectrum and intensity variations in the coastal diatom *Skeletonema marinoi*. Environmental Microbiology 19, 611–627. 10.1111/1462-2920.13545

Soja-Woźniak, M., Clementson, L., Wojtasiewicz, B., Baird, M., 2022. Estimation of the global distribution of phytoplankton light absorption from pigment concentrations. J Geophys Res Oceans 127, e2022JC018494. 10.1029/2022JC018494

Sorigué, D., Légeret, B., Cuiné, S., Blangy, S., Moulin, S., Billon, E., Richaud, P., Brugière, S., Couté, Y., Nurizzo, D., Müller, P., Brettel, K., Pignol, D., Arnoux, P., Li-Beisson, Y., Peltier, G., Beisson, F., 2017. An algal photoenzyme converts fatty acids to hydrocarbons. Science 357, 903–907. 10.1126/science.aan6349

Stonik, V., Stonik, I., 2015. Low-Molecular-Weight Metabolites from Diatoms: Structures, Biological Roles and Biosynthesis. Marine Drugs 13, 3672–3709. 10.3390/md13063672

Su, Y., 2019. The effect of different light regimes on pigments in Coscinodiscus granii. Photosynth Res 140, 301–310. 10.1007/s11120-018-0608-7

Svenning, J.B., Vasskog, T., Campbell, K., Bæverud, A.H., Myhre, T.N., Dalheim, L., Forgereau, Z.L., Osanen, J.E., Hansen, E.H., Bernstein, H.C., 2024. Lipidome Plasticity Enables Unusual Photosynthetic Flexibility in Arctic vs. Temperate Diatoms. Marine Drugs 22, 67. 10.3390/md22020067

Tanaka, T., Yoneda, K., Maeda, Y., 2022. Lipid Metabolism in Diatoms, in: Falciatore, A., Mock, T. (Eds.), The Molecular Life of Diatoms. Springer International Publishing, Cham, pp. 493–527. 10.1007/978-3-030-92499-7_18

Underwood GJC, Barnett, M, 2006. What determines species composition in microphytobenthic biofilms. Royal Netherlands Academy of Arts and Sciences 123–140.

Underwood, G.J.C., Dumbrell, A.J., McGenity, T.J., McKew, B.A., Whitby, C., 2022. The Microbiome of Coastal Sediments, in: Stal, L.J., Cretoiu, M.S. (Eds.), The Marine Microbiome, The Microbiomes of Humans, Animals, Plants, and the Environment. Springer International Publishing, Cham, pp. 479–534. 10.1007/978-3-030-90383-1_12

Valle, K.C., Nymark, M., Aamot, I., Hancke, K., Winge, P., Andresen, K., Johnsen, G., Brembu, T., Bones, A.M., 2014. System responses to equal doses of photosynthetically usable radiation of blue, green, and red Light in the marine diatom *Phaeodactylum tricornutum*. PLoS ONE 9, e114211. 10.1371/journal.pone.0114211

Van Den Dool, H., Dec Kratz, P., 1963. A generalization of the retention index system including linear temperature programmed gas—liquid partition chromatography. Journal of Chromatography A 11, 463–471. 10.1016/S0021-9673(01)80947-X

Van Leeuwe, M.A., Brotas, V., Consalvey, M., Forster, R.M., Gillespie, D., Jesus, B., Roggeveld, J., Gieskes, W.W.C., 2008. Photoacclimation in microphytobenthos and the role of xanthophyll pigments. European Journal of Phycology 43, 123–132. 10.1080/09670260701726119

Wagner, H., Jakob, T., Fanesi, A., Wilhelm, C., 2017. Towards an understanding of the molecular regulation of carbon allocation in diatoms: the interaction of energy and carbon allocation. Phil. Trans. R. Soc. B 372, 20160410. 10.1098/rstb.2016.0410

Walpersdorf, E., Kühl, M., Elberling, B., Andersen, T., Hansen, B., Pejrup, M., Glud, R., 2017. In situ oxygen dynamics and carbon turnover in an intertidal sediment (Skallingen, Denmark). Mar. Ecol. Prog. Ser. 566, 49–65. 10.3354/meps12016

Waring, J., Klenell, M., Bechtold, U., Underwood, G.J.C., Baker, N.R., 2010. Light-induced responses of oxygen photoreduction, reactive oxygen species production and scavenging in two diatom species. J Phycol 46, 1206–1217. 10.1111/j.1529-8817.2010.00919.x

Xu, C., Pi, X., Huang, Y., Han, G., Chen, X., Qin, X., Huang, G., Zhao, S., Yang, Y., Kuang, T., Wang, W., Sui, S.-F., Shen, J.-R., 2020. Structural basis for energy transfer in a huge diatom PSI-FCPI supercomplex. Nat Commun 11, 5081. 10.1038/s41467-020-18867-x

Yamamoto, H., Shikanai, T., 2019. PGR5-Dependent Cyclic Electron Flow Protects Photosystem I under Fluctuating Light at Donor and Acceptor Sides. Plant Physiol. 179, 588–600. 10.1104/pp.18.01343

Zulu, N.N., Zienkiewicz, K., Vollheyde, K., Feussner, I., 2018. Current trends to comprehend lipid metabolism in diatoms. Progress in Lipid Research 70, 1–16. 10.1016/j.plipres.2018.03.001

