## Supplementary figures and tables for "Red and blue light cues drive contrasting remodeling of lipophilic metabolites and photophysiology in natural benthic diatom biofilms"

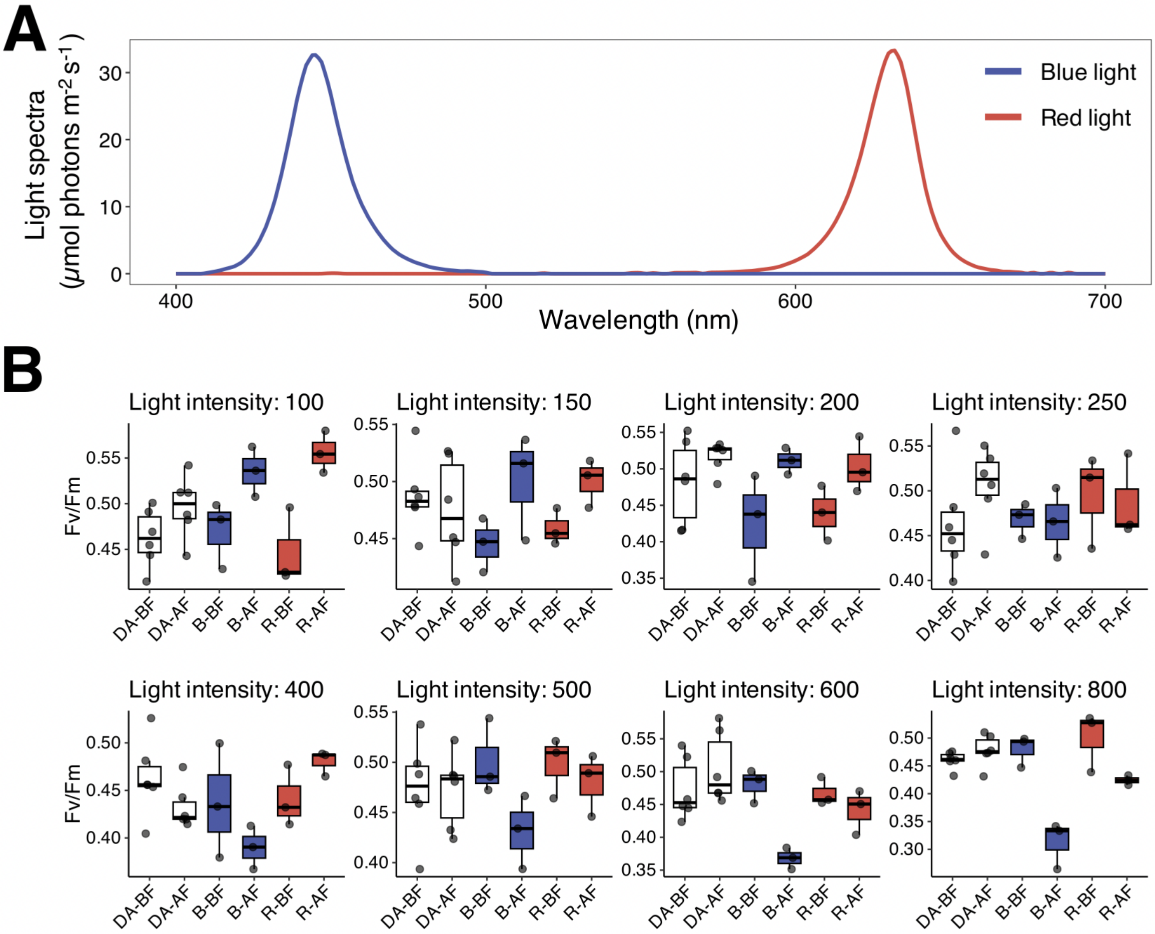


**Figure s1. Incident monochromatic red and blue light spectra and preliminary assessment of Fv/Fm responses**

**A.** Spectral distribution of blue light ranged from 410 to 500 nm with a peak at 450 nm, and red light ranged from 570 to 680 nm with a peak at 640 nm; **B.** Preliminary assessment of photosynthetic responses of diatoms on the day preceding the experiment (14 January 2025). Diatoms were exposed to a range of red and blue light intensities (100–800 µmol photons $m^{-2}$ $s^{-1}$; n = 3 per condition) to identify light levels producing contrasting photosynthetic responses (Fv/Fm) under equivalent incident intensities. Interestingly, the theoretical optimal light range (100–250 µmol photons $m^{-2}$ $s^{-1}$) and the light-induced stress threshold (250–500 µmol photons $m^{-2}$ $s^{-1}$) reported for microphytobenthos from the same sampling site by Doose and Hubas (2024) are consistent with the Fv/Fm responses observed under blue light. Abbreviations: dark-adapted control (including red- and blue-light dark-adapted controls), DA; red light, R; blue light, B; before light exposure, BF; after light exposure and following a 10-minute dark recovery period, AF.

**Table s1. Photosynthetic parameters definitions**

Equations adapted from Consalvey (2005), and Christof Klughammer and Ulrich Schreiber (2008) used to calculate photosynthetic parameters.

| Photosynthetic parameter | Equation | Description |
|---|---|---|
| **α-slope** (arbitrary unit) | $\frac{1}{c}$ | Maximum light use efficiency of photosystem II at low irradiance, derived from the initial slope of the rETR-irradiance (rETR-E) curve. *c : empirical coefficient derived from model fitting.* |
| **Fv/Fm** (unitless) | $\frac{Fm - F0}{Fm}$ | Maximum quantum yield of photosystem II photochemistry under dark-adapted state. *Fm Maximum fluorescence yield of dark-adapted sample; $F_0$ Minimum fluorescence yield of dark-adapted sample.* *Under actinic illumination, this calculation is replaced by Fm'-F/Fm', which corresponds to the effective photosystem II efficiency during subsequent steps of the light curve.* |
| **rETR** (arbitrary unit) | $\frac{Fm' - F}{Fm'} \times \frac{Irradiance}{2}$ | Relative electron transport rate of photosystem II, with the term $rETR_{m\ obs}$ referring to the maximum rETR observed in the light curve measured without extrapolation. *Fm' Maximum fluorescence of illuminated sample; F Fluorescence yield in the light-adapted state measured briefly before application of a Saturation Pulse; Irradiance refers to the incident light µmol photons $m^{-2}$ $s^{-1}$.* |
| **NPQ** (unitless) | $\frac{Fm - Fm'}{Fm'}$ | Non-photochemical quenching, with the term $NPQ^{1000}$ referring to the maximum NPQ value obtained after the final one-minute light curve step at 1000 µmol photons $m^{-2}$ $s^{-1}$, followed by the saturating flash used to determine Fm′. |
| **Y(NPQ)** (unitless) | $\frac{F}{Fm'} - \frac{F}{Fm}$ | Regulated non-photochemical quenching, with the term $Y(NPQ)^{1000}$ referring to the maximum Y(NPQ) value obtained after the final one-minute light curve step at 1000 µmol photons $m^{-2}$ $s^{-1}$, followed by the saturating flash used to determine Fm′. |
| **Y(NO)** (unitless) | $\frac{F}{Fm}$ | Non-regulated non-photochemical quenching, with the term $Y(NO)_{m\ obs}$ referring to the maximum Y(NO) observed within the measured light curve range (no extrapolation). |

**Table s2. Table of identified individual pigments and lipids**

**A.** Table presenting the identified individual pigments and their abbreviations, their retention time during elution (minute), as well as the $\cos^2$ values of each pigment to the first two axes of the MFA analysis; **B.** Table presenting the identified individual lipids, the full set of data associated with these metabolites, as well as the $\cos^2$ values of each lipid to the first two axes of the MFA analysis. *Additional information on identified metabolites is provided in the supplementary .xlsx file available here: [https://github.com/adesparmet/Red-blue-lipophilic-metabolites](https://github.com/adesparmet/Red-blue-lipophilic-metabolites).*

**A**

| Pigment | Retention time (min) | $\cos^2$ PC1 | $\cos^2$ PC2 |
|---|---|---|---|
| **Chlorophyll c2 (Chlc2)** | 9.850 | 0.146 | 0.388 |
| **Fucoxanthin (F)** | 11.870 | 0.013 | 0.147 |
| **Unknown Carotenoid 1 (Uc1)** | 12.690 | 0.129 | 0.099 |
| **Unknown Carotenoid 2 (Uc2)** | 13.100 | 0.609 | 0.006 |
| **Unknown Carotenoid 10 (Uc10)** | 13.360 | 0.614 | 0.029 |
| **Pheophorbide *a* (Pda)** | 13.800 | 0.804 | 0.021 |
| **Unknown Carotenoid 3 (Uc3)** | 14.260 | 0.090 | 0.300 |
| **Unknown Carotenoid 4 (Uc4)** | 14.690 | 0.015 | 0.656 |
| **Diadinoxanthin (Dd)** | 14.940 | 0.116 | 0.266 |
| **Unknown Carotenoid 5 (Uc5)** | 15.180 | 0.095 | 0.216 |
| **Unknown Carotenoid 6 (Uc6)** | 15.270 | 0.145 | 0.180 |
| **Unknown Carotenoid 7 (Uc7)** | 15.570 | 0.030 | 0.035 |
| **Unknown Carotenoid 8 (Uc8)** | 15.660 | 0.455 | 0.161 |
| **Diatoxanthin (Dt)** | 16.740 | 0.106 | 0.316 |
| **Unknown Carotenoid 9 (Uc9)** | 17.250 | 0.428 | 0.191 |
| **Chlorophyll *a* like 1 (Chla-l1)** | 19.000 | 0.046 | 0.053 |
| **Chlorophyll *a* like 2 (Chla-l2)** | 19.460 | 0.031 | 0.002 |
| **Chlorophyll *a* like 3 (Chla-l3)** | 21.250 | 0.013 | 0.440 |
| **Chlorophyll *a* like 4 (Chla-l4)** | 21.700 | 0.006 | 0.425 |
| **Chlorophyll *a* allomer (Chla-allomer)** | 22.160 | 0.867 | 0.007 |
| **Chlorophyll *a* like 5 (Chla-l5)** | 22.500 | 0.812 | 0.002 |
| **Chlorophyll *a* (Chla)** | 22.800 | 0.191 | 0.308 |
| **Chlorophyll *a* epimer (Chla-epimer)** | 23.330 | 0.193 | 0.322 |
| **Chlorophyll *a* like 6 (Chla-l6)** | 23.960 | 0.028 | 0.383 |
| **Chlorophyll *a* like 7 (Chla-l7)** | 24.280 | 0.306 | 0.168 |
| **Chlorophyll *a* like 8 (Chla-l8)** | 24.500 | 0.336 | 0.055 |
| **Chlorophyll *a* like 9 (Chla-l9)** | 25.100 | 0.133 | 0.265 |
| **Chlorophyll *a* like 10 (Chla-l10)** | 25.240 | 0.002 | 0.275 |
| **Pheophytin *a* like 1 (Phal1)** | 25.690 | 0.581 | 0.086 |
| **Pheophytin *a* like 2 (Phal2)** | 25.950 | 0.387 | 0.006 |
| **Pheophytin *a* (Pha)** | 26.160 | 0.338 | 0.077 |
| **Pheophytin *a* like 3 (Phal3)** | 26.420 | 0.350 | 0.014 |
| **$\beta$-$\varepsilon$-carotene ($\beta\varepsilon$)** | 26.790 | 0.003 | 0.298 |
| **$\beta$-$\beta$-carotene ($\beta\beta$)** | 27.120 | 0.194 | 0.020 |
| **Pyropheophytin *a* (Pya)** | 27.370 | 0.239 | 0.120 |

**B**

| Compound | Name | InChIKey | RI calculated | RI NIST | $R_{match}$ NIST | Probability NIST | $cos^2$ PC1 | $cos^2$ PC2 |
|---|---|---|---|---|---|---|---|---|
| **C1** | Isotridecanoic acid (13:0 iso) | SIOLDWZBFABPJU-UHFFFAOYSA-N | 1601.165 | 1588 | 884 | 17.800 | 0.001 | 0.277 |
| **C2** | Tridecanoic acid (13:0) | SZHOJFHSIKHZHA-UHFFFAOYSA-N | 1634.869 | 1624 | 929 | 72.300 | 0.019 | 0.380 |
| **C3** | Isotetradecanoic acid (14:0 iso) | YYVJAABUJYRQJO-UHFFFAOYSA-N | 1699.061 | 1686 | 861 | 44.000 | 0.028 | 0.397 |
| **C4** | Myristoleic Acid (14:1n-5 cis) | YWWVWXASSLXJHU-WAYWQWQTSA-N | 1714.691 | 1715 | 907 | 54.400 | 0.010 | 0.391 |
| **C5** | Tetradecanoic acid (14:0) | TUNFSRHWOTWDNC-UHFFFAOYSA-N | 1734.863 | 1725 | 952 | 71.800 | 0.002 | 0.370 |
| **C6** | 9-Methyltetradecanoic acid (9-Me-15:0) | KXCURTJMVUYHPS-UHFFFAOYSA-N | 1777.546 | 1775 | 765 | 0.440 | 0.104 | 0.018 |
| **C7** | 4,8,12-Trimethyltridecanoic acid (4,8,12-Me-16:0) | FUYCAQNCWDAOLQ-UHFFFAOYSA-N | 1784.248 | 1686 | 704 | 17.300 | 0.144 | 0.142 |
| **C8** | 13-Methyltetradecanoic Acid (15:0 iso) | ZOCYQVNGROEVLU-UHFFFAOYSA-N | 1798.179 | 1779 | 819 | 11.100 | 0.052 | 0.499 |
| **C9** | Methyl 2-(ethoxycarbonylamino)-3-phenylpropanoate | JDBGCNRESYAUON-UHFFFAOYSA-N | 1800.416 | 1783 | 870 | 76.000 | 0.118 | 0.009 |
| **C10** | (Z)6,(Z)9-Pentadecadien-1-ol | AVCYGYFZVWNDGB-HZJYTTRNSA-N | 1805.794 | 1771 | 843 | 2.540 | 0.327 | 0.017 |
| **C11** | 12-Methyltetradecanoate | XKLJLHAPJBUBNL-UHFFFAOYSA-M | 1806.121 | 1788 | 810 | 27.000 | 0.138 | 0.010 |
| **C12** | cis,cis,cis-7,10,13-Hexadecatrienal (16:3n-3) | USMAGXMGZXKLHT-PDBXOOCHSA-N | 1811.172 | 1824 | 894 | 2.590 | 0.012 | 0.422 |
| **C13** | 10Z-pentadecenoic acid (15:1n-5 cis) | APXSAEQXOXTDAM-WAYWQWQTSA-N | 1818.170 | 1817 | 945 | 58.300 | 0.048 | 0.517 |
| **C14** | (Z,E)-9,12-Tetradecadienyl acetate | ZZGJZGSVLNSDPG-FDTUMDBZSA-N | 1824.986 | 1811 | 719 | 1.100 | 0.090 | 0.422 |
| **C15** | Pentadecanoic acid (15:0) | WQEPLUUGTLDZJY-UHFFFAOYSA-N | 1829.972 | 1820 | 955 | 78.400 | 0.030 | 0.042 |
| **C16** | | | 1831.612 | | | | 0.595 | 0.006 |
| **C17** | Neophytadiene | NIDGCIPAMWNKOA-UHFFFAOYSA-N | 1837.547 | 1837 | 897 | 39.800 | 0.466 | 0.097 |
| **C18** | 3-Octadecyne | UQMLWEGZEBCVLH-UHFFFAOYSA-N | 1847.038 | 1828 | 765 | 1.400 | 0.068 | 0.018 |
| **C19** | 6,10,14-Trimethyl-2-pentadecanone | WHWDWIHXSPCOKZ-UHFFFAOYSA-N | 1855.451 | 1844 | 810 | 45.400 | 0.411 | 0.092 |
| **C20** | (E)-11-Hexadecen-1-ol | RHVMNRHQWXIJIS-AATRIKPKSA-N | 1861.976 | 1867 | 760 | 0.160 | 0.420 | 0.001 |
| **C21** | 3-Octadecyne | UQMLWEGZEBCVLH-UHFFFAOYSA-N | 1871.814 | 1828 | 828 | 0.550 | 0.000 | 0.246 |
| **C22** | 9-octadecyne | NKRBWIXXEQOWRY-UHFFFAOYSA-N | 1878.105 | 1828 | 715 | 2.760 | 0.430 | 0.080 |
| **C23** | 7-Octadecyne,2-methyl- | WQMKDHREUPHQQE-UHFFFAOYSA-N | 1891.196 | 1863 | 843 | 0.380 | 0.478 | 0.005 |
| **C24** | (6Z,9Z,12Z)-6,9,12,15-Hexadecatetraenoic acid (16:4) | OKBAWLHHZWRDBE-YSTUJMKBSA-N | 1897.976 | 1863 | 833 | 20.000 | 0.566 | 0.119 |
| **C25** | (6Z,9Z,12Z)-6,9,12-Hexadecatrienoic acid (16:3n-4) | SBEXLJDCZSJDLW-YSTUJMKBSA-N | 1905.059 | 1871 | 805 | 29.800 | 0.127 | 0.134 |
| **C26** | 9-Hexadecenoic acid (16:1n-7 cis) | SECPZKHBENQXJG-FPLPWBNLSA-N | 1916.900 | 1898 | 916 | 22.400 | 0.009 | 0.175 |
| **C27** | (Z)-hexadec-11-enoic acid (16:1n-5 cis) | JGMYDQCXGIMHLL-WAYWQWQTSA-N | 1918.296 | 1913 | 705 | 21.200 | 0.434 | 0.004 |
| **C28** | 7-Hexadecenoic acid (16:1n-9) | PJHOFUXBXJNUAC-UHFFFAOYSA-N | 1924.853 | 1900 | 628 | 6.900 | 0.172 | 0.006 |
| **C29** | Tetradecanoic acid, 10,13-dimethyl-, methyl ester | YMXLTDXXQGMDPS-UHFFFAOYSA-N | 1931.540 | 1926 | 861 | 0.060 | 0.606 | 0.109 |
| **C30** | Hexadecanoic acid (16:0) | IPCSVZSSVZVIGE-UHFFFAOYSA-N | 1939.066 | 1926 | 943 | 72.400 | 0.375 | 0.000 |
| **C31** | | | 1952.936 | | | | 0.313 | 0.000 |
| **C32** | 3-Ureidopropionic acid, N-dimethylaminomethylene-, butyl ester | MQDGDRCOOTULSD-UKTHLTGXSA-N | 1956.710 | 1974 | 748 | 1.060 | 0.192 | 0.003 |
| **C33** | (Z)-7-Hexadecenoic acid (16:1n-9 cis) | PJHOFUXBXJNUAC-KTKRTIGZSA-N | 1960.877 | 1961 | 654 | 0.460 | 0.401 | 0.029 |
| **C34** | Tridecanedioic acid | DXNCZXXFRKPEPY-UHFFFAOYSA-N | 1963.953 | 1960 | 735 | 75.500 | 0.066 | 0.016 |
| **C35** | 3-Eicosyne | AQBXRWJMJJJCLI-UHFFFAOYSA-N | 1970.534 | 1988 | 669 | 4.950 | 0.188 | 0.124 |
| **C36** | 10-Methylhexadecanoic acid | ITFOCFQAVPXNEV-UHFFFAOYSA-N | 1977.339 | 1914 | 598 | 3.510 | 0.286 | 0.002 |
| **C37** | Anteisoheptadecanoic acid (17:0 anteiso) | FXUKWLSZZHVEJD-UHFFFAOYSA-N | 1984.682 | 1984 | 677 | 7.900 | 0.255 | 0.058 |
| **C38** | Isoheptadecanoic acid (17:0 iso) | IIUXHTGBZYEGHI-UHFFFAOYSA-N | 1996.447 | 1990 | 734 | 32.600 | 0.359 | 0.069 |
| **C39** | Ethyl palmitate | XIRNKXNNONJFQO-UHFFFAOYSA-N | 2005.196 | 1993 | 554 | 30.500 | 0.220 | 0.007 |
| **C40** | Eicosane | CBFCDTFDPHXCNY-UHFFFAOYSA-N | 2010.277 | 2000 | 832 | 2.410 | 0.007 | 0.069 |
| **C41** | Heptadecenoic Acid (17:1n-7 cis) | GDTXICBNEOEPAZ-FPLPWBNLSA-N | 2015.671 | 2016 | 803 | 34.300 | 0.003 | 0.004 |
| **C42** | 3,7-Dimethyloct-6-en-1-yl nonanoate | ULWMBBYWQZNAMT-UHFFFAOYSA-N | 2023.252 | 2016 | 629 | 0.650 | 0.113 | 0.203 |
| **C43** | Linolelaidic acid (18:2n-6) | OYHQOLUKZRVURQ-AVQMFFATSA-N | 2021.447 | 2183 | 806 | 6.370 | 0.485 | 0.019 |
| **C44** | Heptadecanoic Acid (17:0) | KEMQGTRYUADPNZ-UHFFFAOYSA-N | 2036.512 | 2028 | 888 | 65.300 | 0.331 | 0.102 |
| **C45** | | | 2034.217 | | | | 0.010 | 0.229 |
| **C46** | Oleyl Alcohol | ALSTYHKOOCGGFT-KTKRTIGZSA-N | 2045.682 | 2006 | 576 | 0.540 | 0.183 | 0.016 |
| **C47** | Methyl octadeca-9-yn-11-trans-enoate | QMZUILXAAOEXNR-CMDGGOBGSA-N | 2047.316 | 2103 | 725 | 1.650 | 0.073 | 0.002 |
| **C48** | 9,12,15-Octadecatrien-1-ol, (Z,Z,Z)- | IKYKEVDKGZYRMQ-PDBXOOCHSA-N | 2056.983 | 2058 | 802 | 1.340 | 0.379 | 0.103 |
| **C49** | 17-Octadecen-14-yn-1-ol | GFIMPQUXRWWOQN-UHFFFAOYSA-N | 2062.994 | 2061 | 635 | 0.150 | 0.482 | 0.081 |
| **C50** | Phytol | BOTWFXYSPFMFNR-PYDDKJGSSA-N | 2076.150 | 2045 | 769 | 4.080 | 0.099 | 0.014 |

| Compound | Name | InChIKey | RI calculated | RI NIST | $R_{match}$ NIST | Probability NIST | $cos^2$ PC1 | $cos^2$ PC2 |
|---|---|---|---|---|---|---|---|---|
| **C51** | γ-Linolenic acid (18:3n-6 GLA) | VZCCETWTMQHEPK-QNEBEIHSSA-N | 2087.940 | 2101 | 810 | 19.600 | 0.076 | 0.013 |
| **C52** | 2Z-heptadecenoic acid (17:1n15 cis) | GEHPRJRWZDWFBJ-NXVVXOECSA-N | 2084.049 | 2073 | 656 | 1.580 | 0.378 | 0.032 |
| **C53** | Stearidonic Acid (18:4n-3 SDA) | JIWBIWFOSCKQMA-LTKCOYKYSA-N | 2094.714 | 2088 | 875 | 50.300 | 0.645 | 0.001 |
| **C54** | Linolenic Acid (18:3n-3 ALA) | DTOSIQBPPRVQHS-PDBXOOCHSA-N | 2103.924 | 2098 | 834 | 16.700 | 0.036 | 0.012 |
| **C55** | Heneicosane | FNAZRRHPUDJQCJ-UHFFFAOYSA-N | 2108.366 | 2100 | 837 | 5.720 | 0.160 | 0.009 |
| **C56** | Petroselinic Acid (18:1n-12 cis) | CNVZJPUDSLNTQU-SEYXRHQNSA-N | 2110.519 | 2105 | 875 | 9.130 | 0.025 | 0.000 |
| **C57** | cis-13-Octadecenoic acid (18:1n-5 cis) | BDLLSHRIFPDGQB-WAYWQWQTSA-N | 2116.217 | 2126 | 883 | 6.440 | 0.207 | 0.015 |
| **C58** | | | 2128.744 | | | | 0.169 | 0.008 |
| **C59** | | | 2128.259 | | | | 0.301 | 0.056 |
| **C60** | Stearic Acid (18:0) | QIQXTHQIDYTFRH-UHFFFAOYSA-N | 2138.644 | 2128 | 891 | 60.100 | 0.000 | 0.115 |
| **C61** | γ-Linolenic acid (18:3n-6 GLA) | VZCCETWTMQHEPK-QNEBEIHSSA-N | 2177.525 | 2144 | 765 | 0.370 | 0.004 | 0.227 |
| **C62** | Ethyl linolenate | JYYFMIOPGOFNPK-AGRJPVHOSA-N | 2182.041 | 2169 | 539 | 0.580 | 0.002 | 0.020 |
| **C63** | 2-Ethylbutyric acid, pentadecyl ester | GBIMMEKJLFMFMU-UHFFFAOYSA-N | 2189.833 | 2190 | 645 | 4.030 | 0.067 | 0.029 |
| **C64** | Dihomo-γ-Linolenic Acid (20:3n-6 DGLA) | HOBAELRKJCKHQD-QNEBEIHSSA-N | 2193.783 | 2300 | 740 | 5.930 | 0.027 | 0.001 |
| **C65** | Icosapent methyl (20:5n-3 EPA methyl) | QWDCYFDDFPWISL-JEBPEJKESA-N | 2199.678 | 2174 | 851 | 14.200 | 0.525 | 0.018 |
| **C66** | Docosane | HOWGUJZVBDQJKV-UHFFFAOYSA-N | 2207.118 | 2200 | 900 | 6.870 | 0.203 | 0.010 |
| **C67** | cis-10-Nonadecenoic acid (19:1n-9 cis) | BBOWBNGUEWHNQZ-KTKRTIGZSA-N | 2212.991 | 2209 | 842 | 13.500 | 0.004 | 0.221 |
| **C68** | Ethyl stearidonate (20:4n-3 ethyl) | RIDOSNBWMUADGT-AFSLFLIVSA-N | 2218.606 | 2209 | 719 | 2.350 | 0.434 | 0.034 |
| **C69** | 10-Nonadecenoic Acid (19:1n-9) | BBOWBNGUEWHNQZ-UHFFFAOYSA-N | 2222.972 | 2212 | 784 | 8.010 | 0.189 | 0.022 |
| **C70** | Nonadecanoic Acid (19:0) | ISYWECDDZWTKFF-UHFFFAOYSA-N | 2236.852 | 2228 | 767 | 34.300 | 0.319 | 0.030 |
| **C71** | Heptadecane, 9-hexyl- | ZUSMXUXZJCOSPV-UHFFFAOYSA-N | 2243.462 | 2243 | 711 | 7.470 | 0.039 | 0.004 |
| **C72** | Arachidonic Acid (20:4n-6 AA) | YZXBAPSDXZZRGB-DOFZRALJSA-N | 2266.822 | 2274 | 882 | 36.900 | 0.480 | 0.068 |
| **C73** | Eicosapentaenoic Acid (20:5n-3 EPA) | JAZBEHYOTPTENJ-JLNKQSITSA-N | 2276.572 | 2282 | 923 | 47.100 | 0.377 | 0.055 |
| **C74** | (all-Z)-8,11,14,17-Eicosatetraenoic acid (20:4n-3) | HQPCSDADVLFHHO-LTKCOYKYSA-N | 2293.125 | 2308 | 857 | 27.200 | 0.203 | 0.155 |
| **C75** | | | 2289.170 | | | | 0.124 | 0.029 |
| **C76** | 5,8,11,14,17-Eicosapentaenoic Acid (20:5n-3 EPA) | JAZBEHYOTPTENJ-UHFFFAOYSA-N | 2300.442 | 2282 | 849 | 20.100 | 0.027 | 0.270 |
| **C77** | Tricosane | FIGVVZUWCLSUEI-UHFFFAOYSA-N | 2306.513 | 2300 | 838 | 1.060 | 0.031 | 0.003 |
| **C78** | cis-11-Eicosenoic acid (20:1n-9 cis) | BITHHVVYSMSWAG-KTKRTIGZSA-N | 2307.439 | 2306 | 732 | 17.800 | 0.197 | 0.013 |
| **C79** | 11-Eicosenoic Acid (20:1n-9 trans) | BITHHVVYSMSWAG-MDZDMXLPSA-N | 2316.566 | 2297 | 861 | 25.900 | 0.186 | 0.123 |
| **C80** | Eicosanoic Acid (20:0) | VKOBVWXKNCXXDE-UHFFFAOYSA-N | 2336.364 | 2329 | 869 | 72.200 | 0.273 | 0.130 |
| **C81** | Methyl dehydroabietate | PGZCJOPTDHWYES-UHFFFAOYSA-N | 2347.077 | 2336 | 810 | 77.700 | 0.002 | 0.037 |
| **C82** | (8E, 11E, 14E)-icosa-8,11,14-trienoic acid (20:3n-6) | HOBAELRKJCKHQD-YHTMAJSVSA-N | 2360.572 | 2346 | 715 | 0.040 | 0.594 | 0.073 |
| **C83** | 2-Methyltricosane | JNHSEDRFFJZMLH-UHFFFAOYSA-N | 2367.965 | 2363 | 822 | 6.470 | 0.101 | 0.039 |
| **C84** | 3-Methyltricosane | WCAGWYVPTZQWEN-UHFFFAOYSA-N | 2376.879 | 2372 | 628 | 1.370 | 0.014 | 0.068 |
| **C85** | Tetracosane | POOSGDOYLQNASK-UHFFFAOYSA-N | 2405.827 | 2400 | 937 | 14.000 | 0.186 | 0.009 |
| **C86** | Methyl 10-(2-octylcyclopropyl)decanoate | MOTIPGVOEIQBHG-UHFFFAOYSA-N | 2415.424 | 2339 | 841 | 15.600 | 0.305 | 0.059 |
| **C87** | Heneicosanoic Acid (21:0) | CKDDRHZIAZRDBW-UHFFFAOYSA-N | 2443.886 | 2429 | 779 | 30.400 | 0.001 | 0.025 |
| **C88** | Docosahexaenoic (22:6n-3 DHA) | MBMBGCFOFBJSGT-KUBAVDMBSA-N | 2468.970 | 2471 | 867 | 37.200 | 0.576 | 0.012 |
| **C89** | Adrenic Acid (22:4n-6 AdA) | TWSWSIQAPQLDBP-DOFZRALJSA-N | 2481.275 | 2481 | 760 | 0.500 | 0.443 | 0.070 |
| **C90** | Pentacosane | YKNWIILGEFFOPE-UHFFFAOYSA-N | 2505.957 | 2500 | 885 | 5.990 | 0.068 | 0.049 |
| **C91** | Erucic Acid (22:1n-9 cis) | DPUOLQHDNGRHBS-KTKRTIGZSA-N | 2511.388 | 2508 | 900 | 56.900 | 0.446 | 0.015 |
| **C92** | Benzene, 1,1'-[2-methyl-2-(phenylthio)cyclopropylidene]bis- | DKKGXIJGZVZJRU-UHFFFAOYSA-N | 2577.538 | 2601 | 654 | 2.400 | 0.003 | 0.002 |
| **C93** | 1-Cycloheptene-3-carboxylic acid, 5-(diphenylmethylene)-, methyl ester | IQTDAOSBRWQHNJ-UHFFFAOYSA-N | 2559.230 | 2557 | 642 | 5.570 | 0.001 | 0.055 |
| **C94** | | | 2544.965 | | | | 0.004 | 0.103 |
| **C95** | 3-Methylpentacosane | VNSJCJLDAGMPAO-UHFFFAOYSA-N | 2571.986 | 2573 | 722 | 0.960 | 0.108 | 0.047 |
| **C96** | Hexacosane | HMSWAIKSFDFLKN-UHFFFAOYSA-N | 2601.778 | 2600 | 908 | 8.650 | 0.025 | 0.014 |
| **C97** | 22-Tricosenoic Acid (23:1n-1) | YGTSVJQQDISEHZ-UHFFFAOYSA-N | 2601.926 | 2654 | 695 | 6.230 | 0.000 | 0.107 |
| **C98** | Tetracosanoic Acid (24:0) | QZZGJDVWLFXDLK-UHFFFAOYSA-N | | 2728 | 770 | 67.100 | 0.138 | 0.004 |

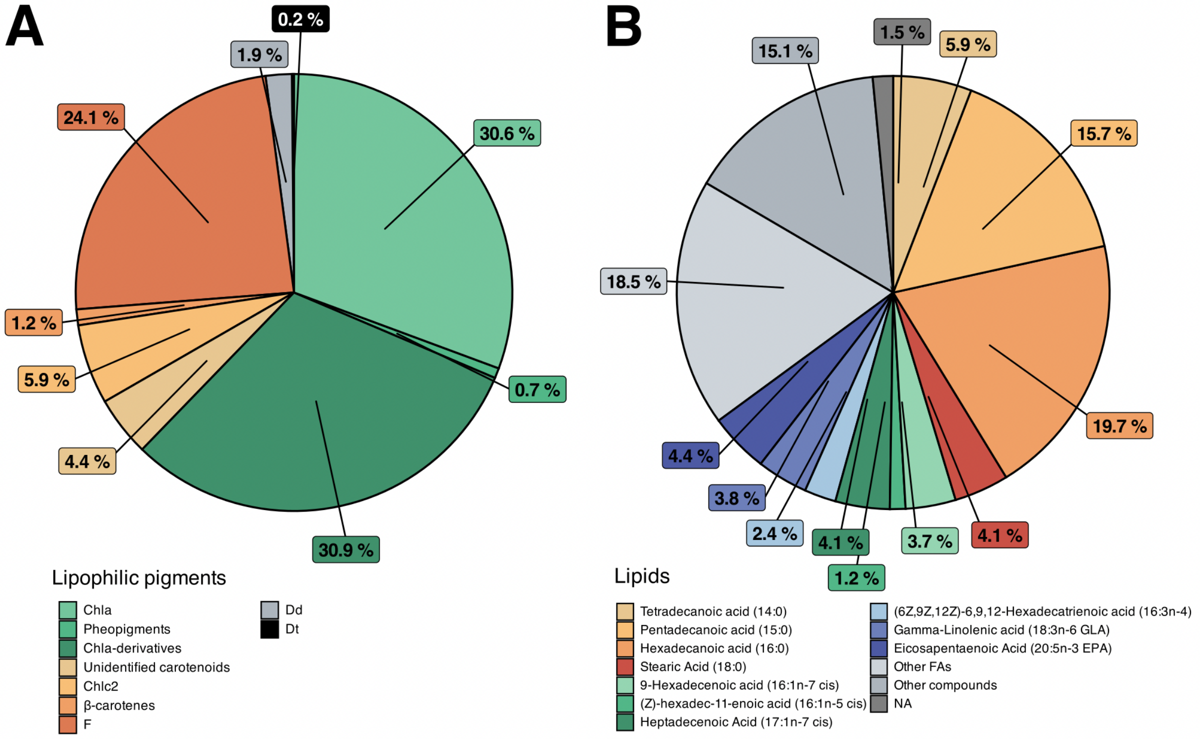


**Figure s2. Average lipophilic metabolite composition**

**A.** Average lipophilic pigment composition (%) across samples (n = 30), highlighting the dominance of chlorophylls, xanthophylls and other carotenoids (pigments abbreviations in Table s2); **B.** Average lipid composition (%) across samples (n = 25), highlighting the dominance of fatty acids. The “Other FAs” category includes fatty acids with < 1% relative abundance. The “Other compounds” category includes the combined relative proportions of alkanes, alkynes, terpenes, amino acids. The “NA” category includes unassigned metabolites.

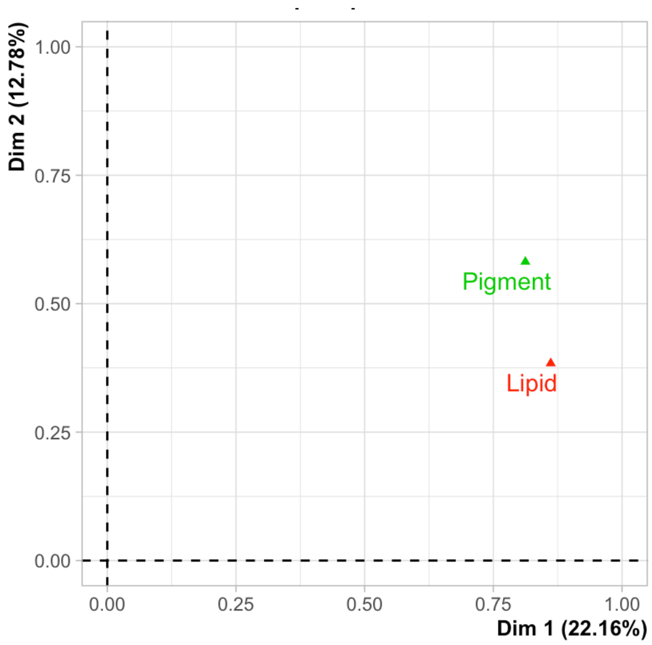


**Figure s3. Representation of the pigment and lipid groups in the MFA**

Contributions of the pigment and lipid groups to the first two dimensions of the MFA. The coordinates of each group on dimensions 1 and 2 indicate their respective associations with the factorial axes and illustrate the extent to which the two data blocks contribute to the structuring of the factorial space.

**Table s3. Mean distances between treatment group centroids**

Pigment and lipid datasets were analyzed separately using Hellinger-transformed compositions (%) to avoid confounding effects of temporal metabolic remodeling, ensuring that the observed structuring and associated Euclidean distances reflected biological variation induced by light treatments. Distances between treatment group centroids were calculated from all PCA dimensions. Light treatments: “Red” dark-adapted control, RC; Red low light, RL; Red high light, RH; “Blue” dark-adapted control, BC; Blue low light, BL; Blue high light, BH.

| Lipophilic metabolites | Comparisons | Distances |
|---|---|---|
| **Lipids** | RC vs RL | 9.71 |
| | RC vs RH | 8.99 |
| | RL vs RH | 6.01 |
| | BC vs BL | 9.23 |
| | BC vs BH | 5.63 |
| | BL vs BH | 6.88 |
| **Pigments** | RC vs RL | 4.97 |
| | RC vs RH | 7.44 |
| | RL vs RH | 5.14 |
| | BC vs BL | 4.30 |
| | BC vs BH | 6.67 |
| | BL vs BH | 5.88 |

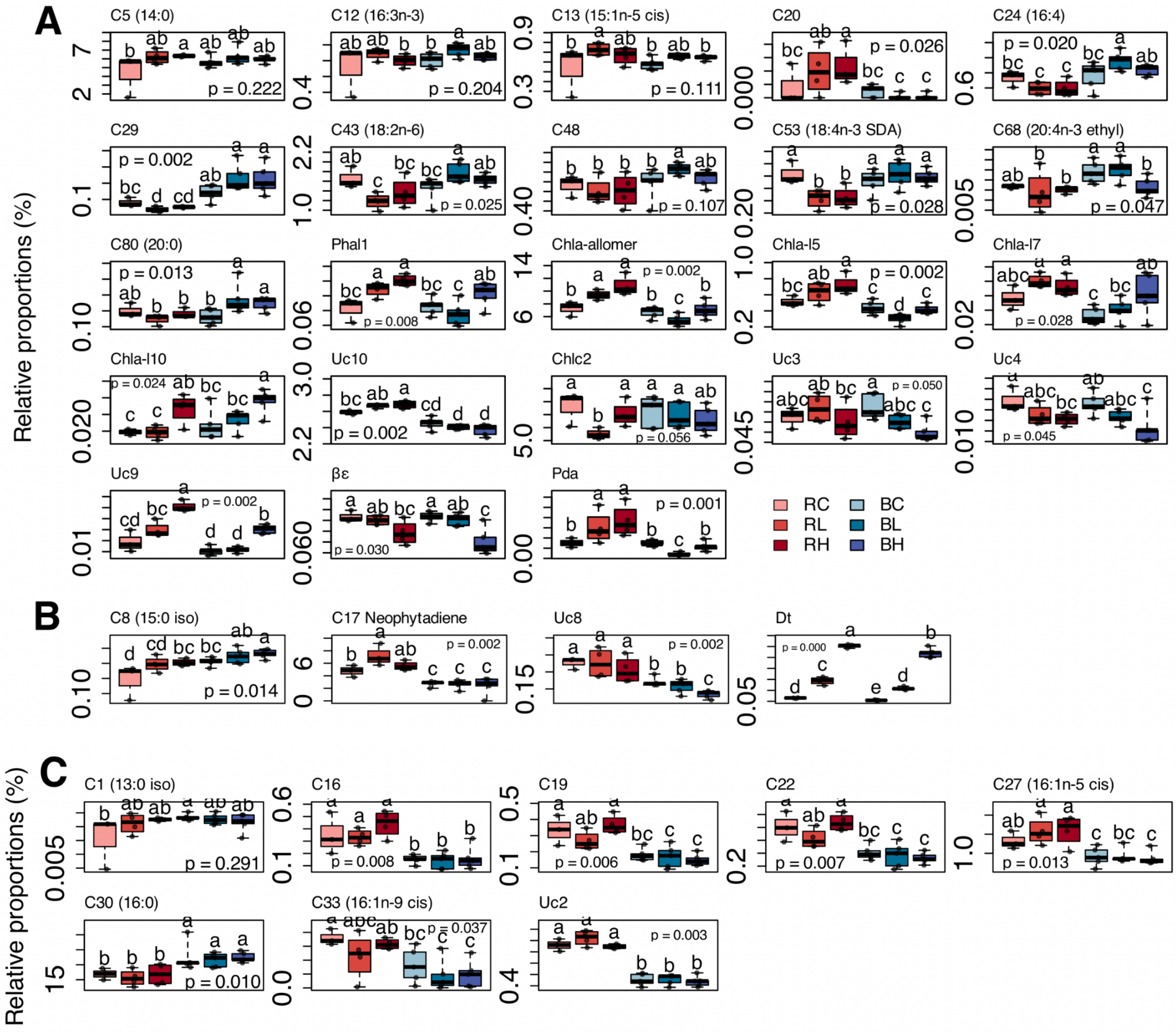
A
C5 (14:0)
C12 (16:3n-3)
C13 (15:1n-5 cis)
C20
C24 (16:4)
C29
C43 (18:2n-6)
C48
C53 (18:4n-3 SDA)
C68 (20:4n-3 ethyl)
C80 (20:0)
Phal1
Chla-allomer
Chla-l5
Chla-l7
Chla-l10
Uc10
Chlc2
Uc3
Uc4
Uc9
βε
Pda
RC
RL
RH
BC
BL
BH
Relative proportions (%)
B
C8 (15:0 iso)
C17 Neophytadiene
Uc8
Dt
C
C1 (13:0 iso)
C16
C19
C22
C27 (16:1n-5 cis)
C30 (16:0)
C33 (16:1n-9 cis)
Uc2
Relative proportions (%)

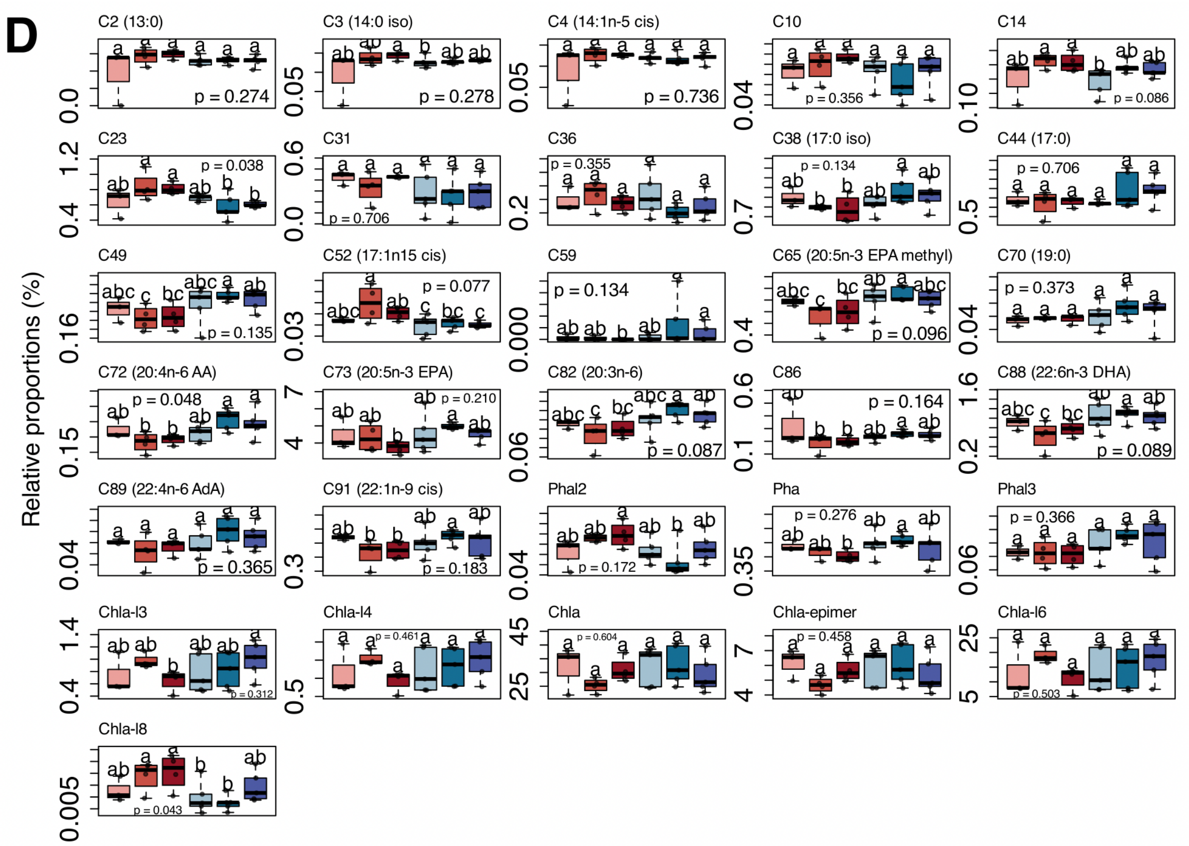


**Figure s4. Individual metabolite variations under light treatments**

Detailed individual dynamics of the top 50% of metabolites most strongly correlated with, at least, one of the first two MFA axes ($cos^2 > 0.27$; n = 66) classified into four response clusters (Fig. 2) according to their variation (%) across light treatments and time (Based on Van der Waerden test results); **A.** Light-responsive metabolites without initial differences in dark-adapted controls; **B.** Metabolites showing both temporal and light-driven variation; **C.** Metabolites affected by temporal variation only; **D.** Stable metabolites with no significant changes across time and treatments. Light treatments: “Red” dark-adapted control, RC; Red low light, RL; Red high light, RH; “Blue” dark-adapted control, BC; Blue low light, BL; Blue high light, BH. Lipids and pigments abbreviations in Table s2.

**Table s4. Xanthophyll concentrations and de-epoxidation states under light treatments**

Average xanthophyll concentrations ($\mu g$ pigments $\mu g^{-1}$ Σ chlorophyll *a* ± SD) and DES values (% ± SD), across light treatments (n = 5). Light treatments precision: “Red” dark-adapted controls, Red control; “Blue” dark-adapted controls, Blue control.

| Treatment | [Diadinoxanthin] | [Diatoxanthin] | [Diadinoxanthin + Diatoxanthin] | DES% |
|---|---|---|---|---|
| **Red control** | 3.24e-02 (1.1e-03) | 1.01e-03 (6.0e-05) | 3.34e-02 (1.1e-03) | 3.02 (0.2) |
| **Red low light** | 3.21e-02 (6.3e-04) | 2.34e-03 (2.5e-04) | 3.44e-02 (4.1e-04) | 6.82 (0.8) |
| **Red high light** | 2.79e-02 (4.7e-04) | 4.87e-03 (1.0e-04) | 3.28e-02 (4.6e-04) | 14.87 (0.4) |
| **Blue control** | 3.35e-02 (8.7e-04) | 8.83e-04 (3.4e-05) | 3.43e-02 (8.9e-04) | 2.57 (0.1) |
| **Blue low light** | 3.22e-02 (7.7e-04) | 1.73e-03 (1.2e-04) | 3.39e-02 (8.4e-04) | 5.11 (0.3) |
| **Blue high light** | 2.84e-02 (9.9e-04) | 4.28e-03 (3.7e-04) | 3.27e-02 (1.0e-03) | 13.11 (1.1) |

**Table s5. Light absorption by the benthic diatom assemblage during the treatments**

This table summarizes the average metrics ± SD (n = 5) used to characterize the light treatments. The incident PPFD column reports the raw incident photosynthetic photon flux density delivered to the samples (µmol photons $m^{-2}$ $s^{-1}$). The Qphar column estimates the portion of incident photon flux actually absorbed by the diatom assemblage during the experiment, without correcting for the cellular pigment package effect (µmol photons $m^{-2}$ $s^{-1}$). The $Qphar_{package\text{-}corrected}$ column shows the same data after accounting for the package effect (µmol photons $m^{-2}$ $s^{-1}$). $P_{abs}$ indicates the total number of photons absorbed by the diatoms over the experimental period ($n_{photons}$), and $E_{abs}$ gives the total quantum energy absorbed by the diatom assemblage during the experiment (J), both calculated from the $Qphar_{package\text{-}corrected}$ values.

| Treatment | PPFD | Qphar | Qphar corrected | Pabs | Eabs |
|---|---|---|---|---|---|
| Blue High Light | 800 | 286.9 (24.7) | 177 (17.1) | 4.6e+20 (4.4e+19) | 203.4 (19.6) |
| Blue Low Light | 200 | 74 (7.6) | 45.9 (5.3) | 1.2e+20 (1.4e+19) | 52.7 (6.1) |
| Red High Light | 800 | 44.7 (3) | 40.8 (2.7) | 1.1e+20 (7.0e+18) | 33.2 (2.2) |
| Red Low Light | 200 | 12.3 (1.4) | 11.2 (1.3) | 2.9e+19 (3.4e+18) | 9.1 (1.1) |

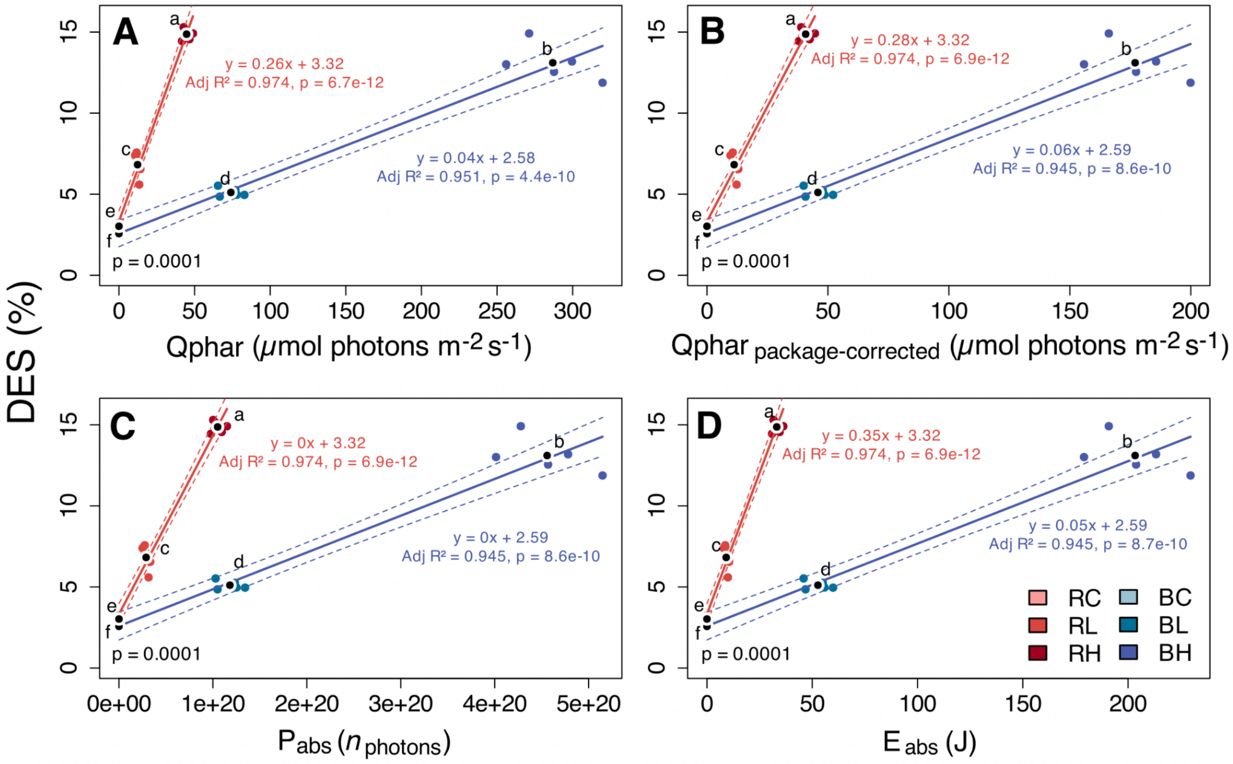


**Figure s5. Linear regression models of DES evolution as a function of Qphar estimates and Pabs and Eabs conversions**

Simple linear regression models showing DES as a function of **A.** the portion of raw incident photosynthetic photon flux density (PPFD) effectively absorbed by the diatoms (Qphar; µmol photons $m^{-2}$ $s^{-1}$), estimated from pigment concentrations and composition profiles and their corresponding reconstructed absorption spectra (Supplementary Fig. 8); **B.** package-corrected Qphar ($Qphar_{package\text{-}corrected}$; µmol photons $m^{-2}$ $s^{-1}$), accounting for the cellular pigment package effect; **C. and D.** total number of photons absorbed ($P_{abs}$; $n_{photons}$) and total absorbed photon energy ($E_{abs}$; J), respectively, both derived from $Qphar_{package\text{-}corrected}$ values. The black dot represents the mean DES value for each treatment group. Differences between treatments were tested using the Van der Waerden test (n = 5). Light treatments: “Red” dark-adapted control, RC; Red low light, RL; Red high light, RH; “Blue” dark-adapted control, BC; Blue low light, BL; Blue high light, BH.

Compared with the representation based on incident PPFD (Fig. 4B), these estimates provide an alternative perspective by considering the amount of light and energy potentially absorbed by the diatom assemblage, rather than relating xanthophyll-cycle responses solely to incident photon flux. This representation suggests that xanthophyll-cycle response dynamics may be influenced by both spectral quality and light intensity. However, these estimates should be interpreted with caution, as the calculations do not account for potential changes in karyostrophy and associated additional pigment-packing effects. Consequently, some values, particularly under blue high-light conditions, may be overestimated. These estimates and representations should therefore be considered indicative rather than quantitatively accurate.

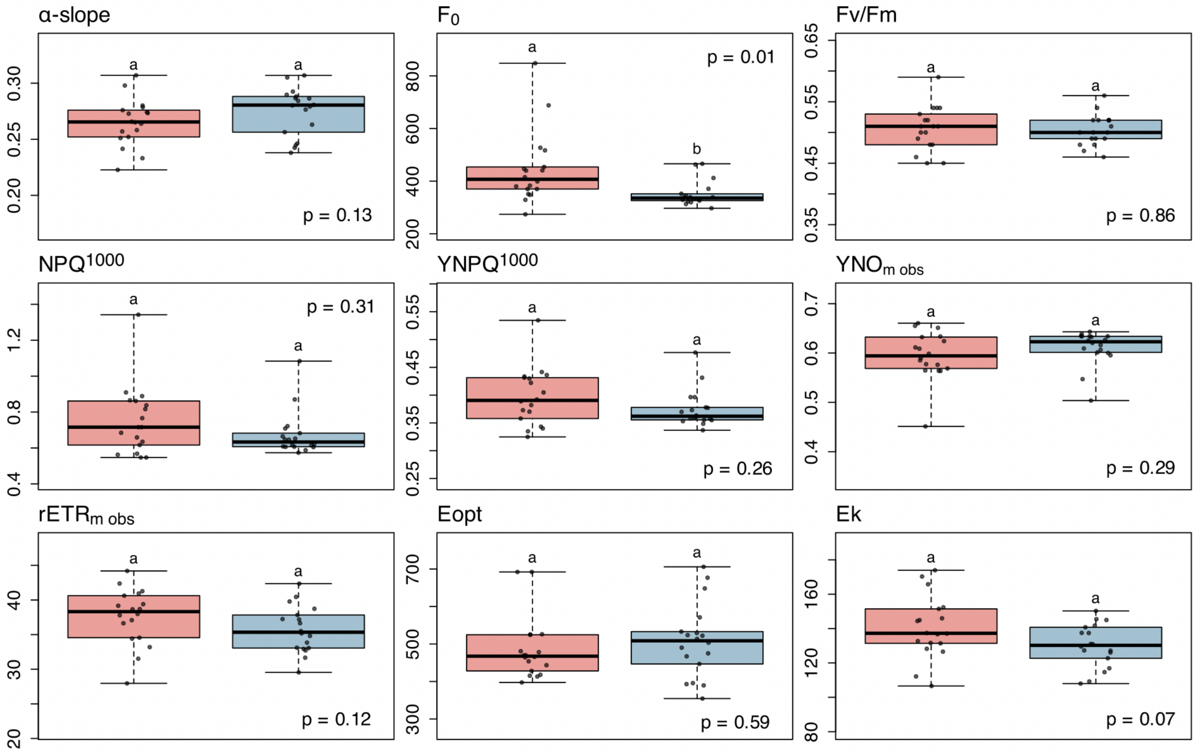


**Figure s6. Dark-adapted photosynthetic parameters throughout the experiment**

Boxplots of all dark-adapted red (morning) and blue (afternoon) samples, including all dark-adapted controls before and after, as well as samples assigned to light treatments but before to their exposure, to assess potential photosynthetic temporal changes without any light cues. α-slope (arbitrary unit) corresponds to the maximum light use efficiency of photosystem II under low irradiances. Fv/Fm (unitless) corresponds to the maximum quantum yield of photosystem II photochemistry under dark-adapted state. $rETR_{m\ obs}$ (arbitrary unit) and $Y(NO)_{m\ obs}$ (unitless) refer to the maximum values observed within the measured light curve range (without extrapolation). $NPQ^{1000}$ (unitless) and $Y(NPQ)^{1000}$ (unitless) correspond to the maximum values measured after the final one-minute light curve step at 1000 µmol photons $m^{-2}$ $s^{-1}$ (without extrapolation). $F_0$ refers to the minimum dark-adapted fluorescence yield (arbitrary unit). Eopt refers to the optimal light parameter for autotrophic cells (µmol photons $m^{-2}$ $s^{-1}$). Ek refers to the light saturation coefficient where the cell begins to activate photoprotective mechanisms (µmol photons $m^{-2}$ $s^{-1}$). All fluorescence measurements were performed on dark-adapted samples.

**Table s6. Table of raw photosynthetic parameters**

Photosynthetic parameter values (mean ± SD; n = 3) before and after light treatments. α-slope (arbitrary unit) corresponds to the maximum light use efficiency of photosystem II under low irradiances. Fv/Fm (unitless) corresponds to the maximum quantum yield of photosystem II photochemistry under dark-adapted state. $rETR_{m\ obs}$ (arbitrary unit) and $Y(NO)_{m\ obs}$ (unitless) refer to the maximum values observed within the measured light curve range (without extrapolation). $NPQ^{1000}$ (unitless) and $Y(NPQ)^{1000}$ (unitless) correspond to the maximum values measured after the final one-minute light curve step at 1000 µmol photons $m^{-2}$ $s^{-1}$ (without extrapolation). $F_0$ refers to the minimum dark-adapted fluorescence yield (arbitrary unit). Eopt refers to the optimal light parameter for autotrophic cells (µmol photons $m^{-2}$ $s^{-1}$). Ek refers to the light saturation coefficient where the cell begins to activate photoprotective mechanisms (µmol photons $m^{-2}$ $s^{-1}$). All fluorescence measurements were performed on dark-adapted samples. Light treatments: "Red high light" dark-adapted control, RHC; "Red low light" dark-adapted control, RLC; Red low light, RL; Red high light, RH; "Blue high light" dark-adapted control, BHC; "Blue low light" dark-adapted control, BLC; Blue low light, BL; Blue high light, BH.

| Treatment | α-slope | Fm/Fv | rETRm obs | NPQ1000 | YNPQ1000 | YNOm obs | F0 | Eopt | Ek |
|---|---|---|---|---|---|---|---|---|---|
| **RH Before** | 0.28 ± 0.05 | 0.51 ± 0.03 | 37.95 ± 4.22 | 0.71 ± 0.16 | 0.39 ± 0.05 | 0.60 ± 0.04 | 403.00 ± 35.17 | 543.93 ± 128.38 | 139.17 ± 31.62 |
| **RH After** | 0.27 ± 0.02 | 0.45 ± 0.02 | 41.76 ± 3.86 | 0.58 ± 0.04 | 0.34 ± 0.02 | 0.68 ± 0.02 | 346.67 ± 7.51 | 1267.96 ± 1240.62 | 159.01 ± 13.18 |
| **RHC Before** | 0.25 ± 0.01 | 0.48 ± 0.03 | 34.87 ± 5.99 | 0.64 ± 0.02 | 0.37 ± 0.01 | 0.63 ± 0.02 | 367.00 ± 14.73 | 453.12 ± 33.27 | 134.45 ± 24.46 |
| **RHC After** | 0.26 ± 0.01 | 0.49 ± 0.05 | 38.11 ± 3.17 | 0.60 ± 0.10 | 0.35 ± 0.04 | 0.63 ± 0.03 | 328.67 ± 54.50 | 521.26 ± 148.24 | 145.75 ± 19.62 |
| **RL Before** | 0.25 ± 0.02 | 0.50 ± 0.03 | 38.88 ± 6.58 | 0.78 ± 0.19 | 0.41 ± 0.05 | 0.60 ± 0.06 | 443.00 ± 90.01 | 468.04 ± 65.24 | 149.68 ± 19.06 |
| **RL After** | 0.26 ± 0.03 | 0.48 ± 0.05 | 37.94 ± 3.73 | 0.69 ± 0.16 | 0.38 ± 0.05 | 0.63 ± 0.06 | 351.33 ± 84.29 | 508.34 ± 14.95 | 140.32 ± 3.08 |
| **RLC Before** | 0.28 ± 0.00 | 0.55 ± 0.04 | 37.81 ± 4.11 | 1.01 ± 0.28 | 0.47 ± 0.06 | 0.53 ± 0.07 | 604.00 ± 214.19 | 428.26 ± 22.23 | 137.11 ± 12.55 |
| **RLC After** | 0.27 ± 0.01 | 0.52 ± 0.02 | 37.86 ± 0.80 | 0.77 ± 0.05 | 0.41 ± 0.02 | 0.59 ± 0.02 | 514.67 ± 150.67 | 504.75 ± 35.00 | 137.07 ± 0.35 |
| **BH Before** | 0.29 ± 0.01 | 0.52 ± 0.01 | 38.17 ± 3.67 | 0.81 ± 0.24 | 0.41 ± 0.06 | 0.57 ± 0.06 | 395.33 ± 59.37 | 557.95 ± 78.72 | 133.02 ± 15.02 |
| **BH After** | 0.16 ± 0.03 | 0.27 ± 0.03 | 33.70 ± 7.18 | 0.31 ± 0.00 | 0.22 ± 0.00 | 0.83 ± 0.01 | 266.33 ± 19.86 | 771.40 ± 87.96 | 222.31 ± 59.78 |
| **BHC Before** | 0.28 ± 0.03 | 0.51 ± 0.05 | 37.41 ± 3.86 | 0.70 ± 0.15 | 0.38 ± 0.05 | 0.60 ± 0.05 | 376.67 ± 77.36 | 564.90 ± 115.15 | 135.43 ± 7.50 |
| **BHC After** | 0.28 ± 0.01 | 0.52 ± 0.02 | 38.29 ± 1.33 | 0.65 ± 0.06 | 0.37 ± 0.02 | 0.61 ± 0.01 | 354.67 ± 51.33 | 464.43 ± 65.88 | 134.57 ± 9.62 |
| **BL Before** | 0.27 ± 0.03 | 0.48 ± 0.02 | 33.52 ± 3.83 | 0.62 ± 0.03 | 0.36 ± 0.01 | 0.62 ± 0.02 | 327.33 ± 10.21 | 549.35 ± 135.66 | 128.40 ± 18.04 |
| **BL After** | 0.28 ± 0.03 | 0.49 ± 0.04 | 43.34 ± 1.44 | 0.74 ± 0.12 | 0.39 ± 0.04 | 0.62 ± 0.02 | 307.00 ± 43.86 | 609.77 ± 110.88 | 154.69 ± 18.67 |
| **BLC Before** | 0.27 ± 0.02 | 0.48 ± 0.01 | 33.89 ± 1.93 | 0.61 ± 0.00 | 0.36 ± 0.00 | 0.64 ± 0.00 | 331.33 ± 6.81 | 484.14 ± 76.78 | 126.11 ± 16.66 |
| **BLC After** | 0.27 ± 0.02 | 0.51 ± 0.02 | 32.92 ± 0.20 | 0.65 ± 0.05 | 0.37 ± 0.02 | 0.62 ± 0.02 | 324.67 ± 24.83 | 425.61 ± 91.96 | 123.01 ± 12.59 |

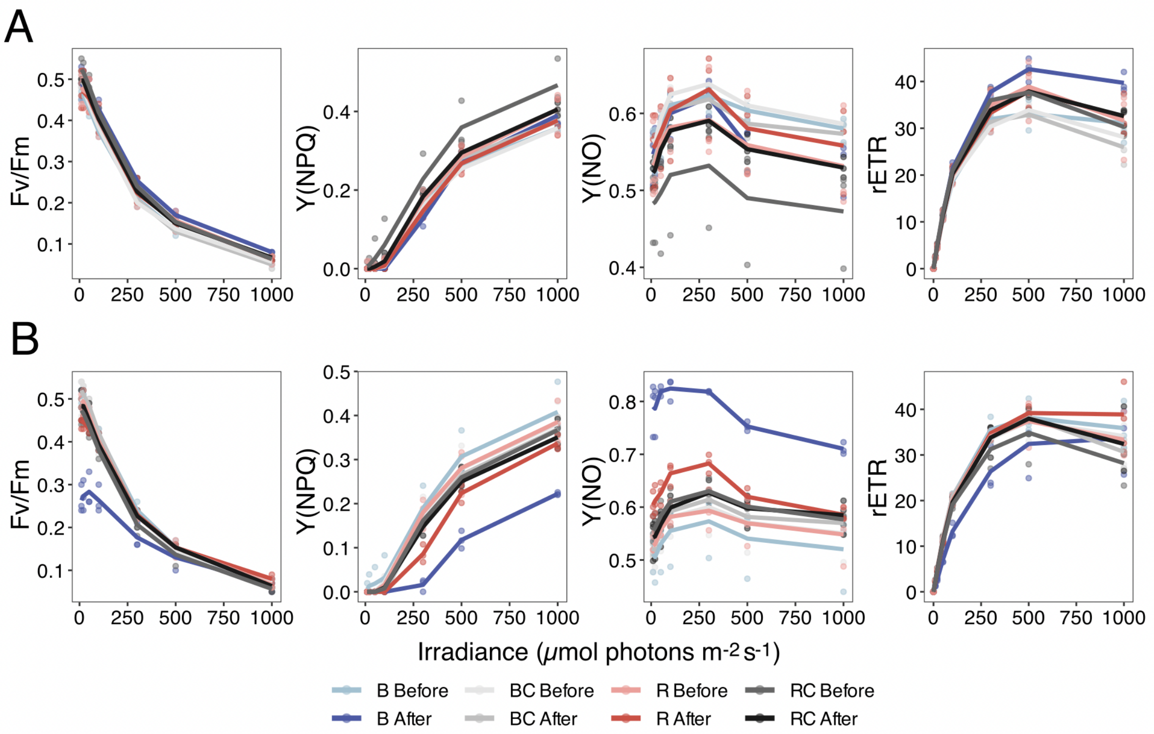


**Figure s7. Light curve responses under light treatments**

Fv/Fm (unitless), Y(NPQ) (unitless), Y(NO) (unitless) and rETR (arbitrary unit) photosynthetic parameters responses; **A.** Rapid light curve of the diatom assemblage before and after low light treatments; **B.** Rapid light curve of the diatom assemblage before and after high light treatments. All photosynthetic parameters were measured on dark-adapted samples. Light treatments (n = 3): “Red” dark-adapted control, RC; “Blue” dark-adapted control, BC; Blue light, B; Red light, R.
