## Supplementary material and method for "Red and blue light cues drive contrasting remodeling of lipophilic metabolites and photophysiology in natural benthic diatom biofilms"

### Supplementary materials and methods

**Quantification of package-corrected photosynthetically usable incident light.** To more accurately quantify the light actually absorbed by the diatom suspension during the treatments, Qphar values were calculated as a refined equivalent of photosynthetically usable radiation (PUR). The pigment profiles of each sample were used to reconstruct their respective absorption spectra, following **Eq.1** adapted from Gilbert et al. (2000) and Soja-Woźniak et al. (2022):

$$a_{microphytobenthos}^{rec,un}\ (\lambda)\ = \sum_{i}^{n}\quad a_i(\lambda) \times C_i \qquad \textbf{(1)}$$

where $a_i(\lambda)$ ($m^2$ $mg^{-1}$) refers to the concentration-specific absorption spectra of pigments provided by Clementson and Wojtasiewicz (2019). $C_i$ (mg $m^{-2}$) corresponds to the pigment concentrations per Petri dish. *n* represents the number of individual microphytobenthic pigments included in the reconstruction of the absorption spectrum of the diatom suspension, with all pigments listed in Supplementary Table 2A being considered.

Although larger diatoms (e.g., *P. strigosum*, *Gyrosigma sp.*) appear to have evolved compensatory mechanisms, self-shading reduces their absorption efficiency per pigment unit, underscoring the need to estimate the package effect alongside Qphar to better approximate the light actually absorbed by the diatom suspension (Supplementary Fig. 8) (Gilbert et al., 2000; Malerba et al., 2018; Soja-Woźniak et al., 2022). A correction for the cellular package effect was estimated from the "typical" absorption spectrum ($a_{microphytobenthos}{}^{rec,un}$; unitless) reconstructed from undisturbed dark-adapted controls, using the equations provided by Soja-Woźniak et al. (2022). This approach quantifies the percentage of light attenuation at each wavelength caused by intracellular pigment packing (i.e., the modulation of light absorption per pigment unit as a function of pigment concentration and cell size). The estimated wavelength-specific attenuation values were then applied to all light-treated samples. The resulting corrected reconstructed absorption spectra ($a_{microphytobenthos}{}^{rec,pac}$; unitless) were subsequently weighted by raw incident light treatments in $Qphar_{package\text{-}corrected}$ calculations, using **Eq.2** based on equation from Gilbert et al. (2000) and Soja-Woźniak et al. (2022):

$$Qphar_{package-corrected}(\lambda)\ =\ Q(\lambda)\ -\ (Q(\lambda) \times\ e^{-a_{microphytobenthos}^{rec,pac}(\lambda)}) \qquad \textbf{(2)}$$

where *Q(λ)* (µmol photons $m^{-2}$ $s^{-1}$) represents the raw incident PPFD available to the diatom suspension. $a_{microphytobenthos}^{rec,pac}$ is the package-corrected absorption spectrum of the sample (unitless).

From these $Qphar_{package\text{-}corrected}$ values, the total number of photons absorbed ($P_{abs}$; $n_{photons}$) by the diatom suspension over the full duration of treatments was calculated using **Eq.3:**

$$P_{abs} = \Sigma_\lambda \left[ Qphar_{package-corrected}(\lambda) \times S \times t \times 10^{-6} \times N_A \right] \quad \textbf{(3)}$$

The photon flux absorbed at each wavelength ($Qphar_{package\text{-}corrected}$; µmol photons $m^{-2}$ $s^{-1}$) was multiplied by the Petri dish surface area (*S*; $m^2$) and the light exposure time (*t*; s) to obtain the total amount of absorbed photons (µmol). This value was converted to mol ($x10^{-6}$) and then to the number of photons by multiplication with Avogadro's constant (~6.022 $x10^{23}$ $mol^{-1}$).

Absorbed photon quantities at each wavelength were converted into energy (joules) and then summed across wavelengths to obtain the total absorbed quantum energy ($E_{abs}$; J), using the wavelength-dependent photons energy ($E_{photon}$; J) derived from Planck's constant (*h*; 6.626 $x10^{-34}$ J s) and the speed of light in vacuum (*c*; 2.998 $x10^{8}$ m $s^{-1}$), as described in **Eqs.4 and 5**:

$$E_{photon}(\lambda) = \frac{h \times c}{\lambda \times 10^{-9}} \quad \textbf{(4)}$$

$$E_{abs} = \Sigma_\lambda \left[ P_{abs}(\lambda) \times E_{photon}(\lambda) \right] \quad \textbf{(5)}$$

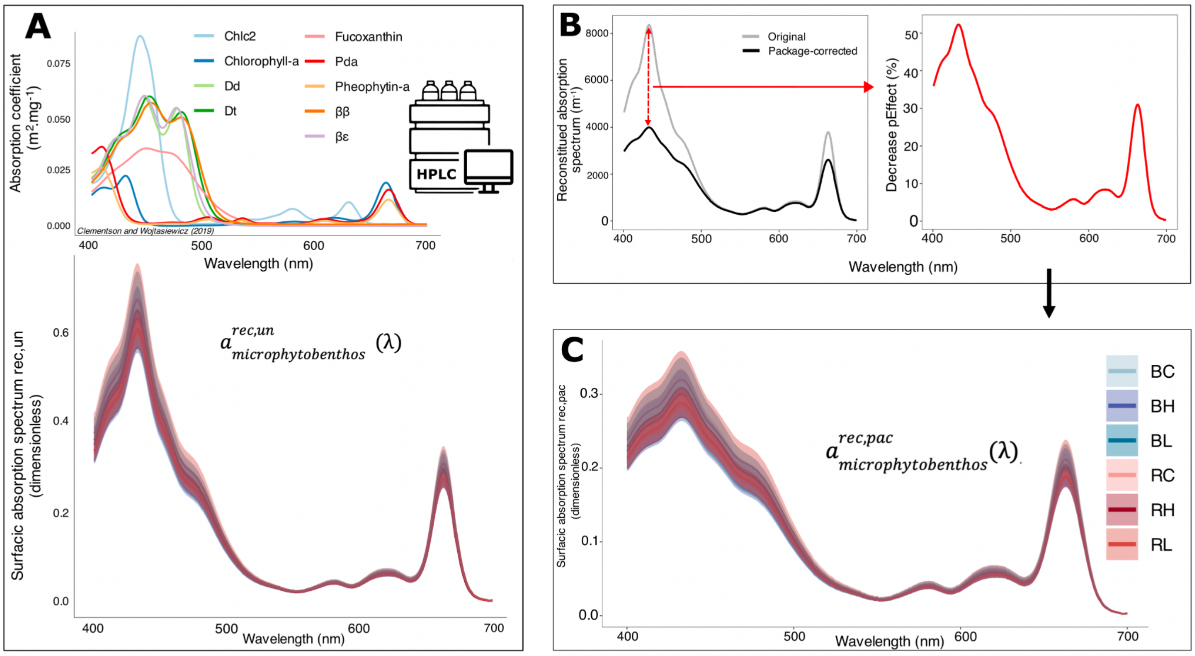


**Figure s8. Absorption spectra reconstruction and package effect correction**

**A.** Pigment-specific absorption coefficients ($m^2$ $mg^{-1}$) used with surfacic pigment concentrations (mg $m^{-2}$) for the reconstruction of the original absorption spectra of the diatom assemblage (reconstructed spectrum uncorrected package-effect; rec,un); **B.** Estimation of the cellular package-effect by calculating the average absorbance loss (%), at each wavelength, from the mean reconstructed dark-adapted controls absorption spectrum; **C.** Package-effect correction of diatom assemblage absorption spectra (reconstructed spectrum package-effect corrected; rec,pac); Application of the mean correction across all replicates to obtain individual absorption spectra corrected with the cellular package-effect. Light treatments: “Red” dark-adapted control, RC; Red low light, RL; Red high light, RH; “Blue” dark-adapted control, BC; Blue low light, BL; Blue high light, BH. Pigments abbreviations in Table s2.

Despite simplifications, particularly by the lack of karyostrophy consideration which strongly increase the packaging effect (Prins et al., 2020; Bastos et al., 2025; Desparmet et al., 2026), the resulting $Qphar_{package\text{-}corrected}$ values indicate that blue light is absorbed approximately 4.2 times more efficiently than red light (Supplementary Table 5), close to the empirical ratio of 4.6 reported in *Phaeodactylum tricornutum* (Valle et al., 2014).
